# Type IV competence pili in *Streptococcus pneumoniae* are highly dynamic structures that retract to promote DNA uptake

**DOI:** 10.1101/2021.01.17.426607

**Authors:** Trinh Lam, Courtney K. Ellison, Ankur B. Dalia, David T. Eddington, Donald A. Morrison

## Abstract

The competence pili of transformable Gram-positive species form a subset of the diverse and widespread class of extracellular filamentous organelles known as type IV pili (T4P). In Gram-negative bacteria, T4P act through dynamic cycles of extension and retraction to carry out diverse activities including attachment, motility, protein secretion, and DNA uptake. It remains unclear whether T4P in Gram-positive species exhibit this same dynamic activity, and their mechanism of action for DNA uptake remains unclear. They are hypothesized to either (1) passively form transient cavities in the cell wall to facilitate DNA passage, (2) act as static adhesins to enrich DNA near the cell surface for subsequent uptake by membrane-embedded transporters, or (3) play an active role in translocating bound DNA via their dynamic activity. Here, using a recently described pilus labeling approach, we demonstrate that pneumococcal competence pili are highly dynamic structures that rapidly extend and retract from the cell surface. By labeling ComGC with bulky adducts, we further demonstrate that pilus retraction is essential for natural transformation. Together, our results indicate that Gram-positive type IV competence pili are dynamic and retractile structures that play an active role in DNA uptake.

**Short summary:** Competent pneumococci kill non-competent cells on contact. Retractable DNA-binding fibers in the class of type IV pili may provide a key tool for retrieving DNA segments from cell wreckage for internalization and recombination.

## INTRODUCTION

Bacteria interact with their environment through various active physical mechanisms. Motility driven by rotating flagella that move the cell body through suspending liquids, for example, allows long-range exploration of liquid milieux, especially when coupled with temporal sensing of solutes of potential value or danger. At closer range, long thin protein appendages of a widespread class known as type IV pili support a different mechanism of directional motility, which operates much like the nautical grapnel or kedge anchor devices, by means of fibers that are alternately extended and retracted [4, 7, 8, 48, 51, 56, 68, 71, 76]. The retractile movement is actuated by one of the strongest biological motors known [46]. Type IV pili also support an array of biological functions not directly related to motility, including surface sensing [24], control of colony shape [5, 34, 65, 74], adhesion to surfaces or host cells [13, 28, 33, 59], and biofilm formation [37, 40], but these functions still depend on the retractile property of the fibers.

A fascinating variant on these type IV pilus-mediated functions is uptake of DNA by competent cells during natural genetic transformation, where the object moved is a DNA molecule, not a cell [1, 29, 32, 41, 64, 69]. In Gram-negative bacteria, some type IV pili, assembled either as microns-long extensions or as short ‘pseudopili’, are absolutely required for such DNA uptake [1, 2, 11, 29, 32, 41, 49, 66, 69, 72]. Among these, the type IV competence pilus of *Vibrio cholerae* mediates DNA uptake by protruding perpendicularly to the cell surface, binding to DNA via its tip, and retracting to translocate bound DNA across the outer membrane into the periplasmic space [25].

Type IV pili are widespread within the Gram-positive bacteria [35], where they have roles in motility and adhesion much like those seen in the Gram-negative groups [63]. Evidence also implicates a type IV pilus-like structure known as the com pilus in horizontal gene transfer in many firmicutes species, where conserved ComG/PilD competence pilus operons are required for DNA uptake, but their mechanism of action remains obscure. The *comG/pilD* operons are most thoroughly characterized in *Bacillus subtilis*, where they encode a short (pseudo)pilus, whose essential role in DNA uptake is thought to depend on reversible movement within the cell wall [9, 16, 15, 17].

All streptococci maintain genes encoding the basic components needed for elaboration of a com pilus that is orthologous to the com competence pseudopilus of *B. subtilis*, and, like *B. subtilis*, regulate their expression to coincide with the development of competence for natural genetic transformation [39]. The *comG* and *pilD* operons, conserved among all competent Gram-positives, comprise eight genes. Structural com pilus subunits include ComGC, the principal subunit called the “major pilin”, and minor pilin subunits designated ComGD, ComGE, ComGF, and ComGG. ComGA is a cytosolic secretion ATPase; ComGB is a polytopic membrane protein; *pilD*, encoding the prepilin peptidase, is unlinked to the *comG* operon, but is coregulated with it during competence induction [39].

Only one Gram-positive competence pilus has thus far been directly observed as an external appendage. Electron microscopy and immunostaining directed against the ComGC pilin of *Streptococcus pneumoniae* (pneumococcus) revealed that competent cultures of this species specifically accumulate ComGC-containing pilus fragments, and that competent cells carry one or two pili with a typical type IV pilus structure, approximately 6 nm thick but 1000 or more nm long, clearly distinct from the paradigmatic *B. subtilis* pseudopilus [44]. A direct role for this pilus in DNA uptake is suggested by the observation that DNA co-purifies with pilus fragments and is found associated lengthwise with pilus fragments via EM analysis [44].

Although type IV competence pili of both Gram-negative and Gram-positive bacteria mediate DNA uptake, there are important differences between the competence pili in the Gram-positive *S. pneumoniae* and those in the Gram-negative *V. cholerae*. First*, V. cholerae* type IV competence pili have separate motor ATPases that mediate their extension and retraction, respectively, whereas pneumococcal and other com pilus loci lack a retraction ATPase [21, 44]. Nonetheless, many pilus systems without a dedicated retraction ATPase still exhibit dynamic retraction [21] and at least one extension ATPase, for the tad pilus, is known to be bifunctional, promoting both pilus extension and retraction [27]. Thus, a major open question about the pneumococcal competence pilus is whether this structure both dynamically extends and retracts. Second, unlike *V. cholerae*, competent Gram-positive bacteria do not translocate DNA across an outer membrane; they might instead use competence pili to bring DNA (via retraction) through the thick cell wall for access to the DNA processing and transport machine. Thus, a second major unanswered question is whether Gram-positive competence pili actively move DNA across the thick peptidoglycan barrier.

Because numerous type IV pili serve functions distinct from motility that nonetheless depend on retraction, our hypothesis was that, despite lacking an apparent retraction ATPase, the streptococcal competence pilus is a retractable organelle, and that, like other type IV pili, its subunits equilibrate, via extension/retraction cycles, with a pool of monomers in the membrane. The pneumococcal competence pilus was previously visualized using a FLAG epitope-directed fluorescent antibody in fixed preparations [44]. However, the bulkiness of such a label moiety is likely to interfere with any retractive pilus activity in living bacteria. To obtain a more dynamic view of the pneumococcal competence pilus, we coupled the less bulky thiol-reactive fluorescent maleimide conjugate Alexa-Fluor 488 C5 maleimide (AF488-mal) directly to cysteine residues that we placed, by targeted mutagenesis, within the ComGC major pilin [26]. We report here that the strategy labels extracellular pili and that these tagged competence pili are highly dynamic, participating in cycles of extrusion and retraction over timescales of seconds. We further provide evidence that this retraction is required for DNA uptake, consistent with the hypothesis that these pili act as fishing lines for DNA.

## RESULTS

### Cys substitution mutants of S. pneumoniae

Pneumococcal type IV competence pili are long filamentous 6.4-nanometer-diameter appendages composed almost entirely of a single repeating pilin subunit, the major pilin ComGC, which lacks native cysteines [44, 54]. To label these structures, cysteine knock-in mutants of ComGC can be generated, which, if the incorporated cysteine is solvent accessible, would allow for subsequent labeling of pili with low-MW fluorescently conjugated maleimide dyes [26]. While this technique has been used successfully to label pili from diverse bacterial species, the first hurdle in this approach is to obtain mutant strains where neither the substituted cysteine nor the maleimide adduct in the major pilin compromises pilus function. Encouraged by the fact that a FLAG epitope placed at the C-terminus of ComGC does not interfere with transformation [44], indicating that other small alterations to the pilin structure might be tolerated as well, we designed and constructed nine distinct pneumococcal Cys-substituted *comGC* mutants, based on analysis of the ComGC subunit to pinpoint likely surface-exposed residues (**Fig. 1 A-B**). All of the new *comGC*-Cys substitution mutants have functional type IV competence pili, as evidenced by detectable rates of natural transformation (**Fig. 1C**). Three transformed at rates indistinguishable from the parent, while the remaining six retained readily detectable but reduced rates of transformation. The substitutions with the highest transformation rates were at serine residues S60 and S66 in the N-terminal alpha 1-C core domain, or at S74, in the adjacent alpha 2 region of the head domain [54]. Thus, although not all Cys replacements are neutral toward transformation, at least three innocuous positions are available for placement of a Cys thiol target for maleimide labeling. Most results reported here were obtained with SAD1671, carrying the ComGC^S66C^ substitution.

**Figure 1.**
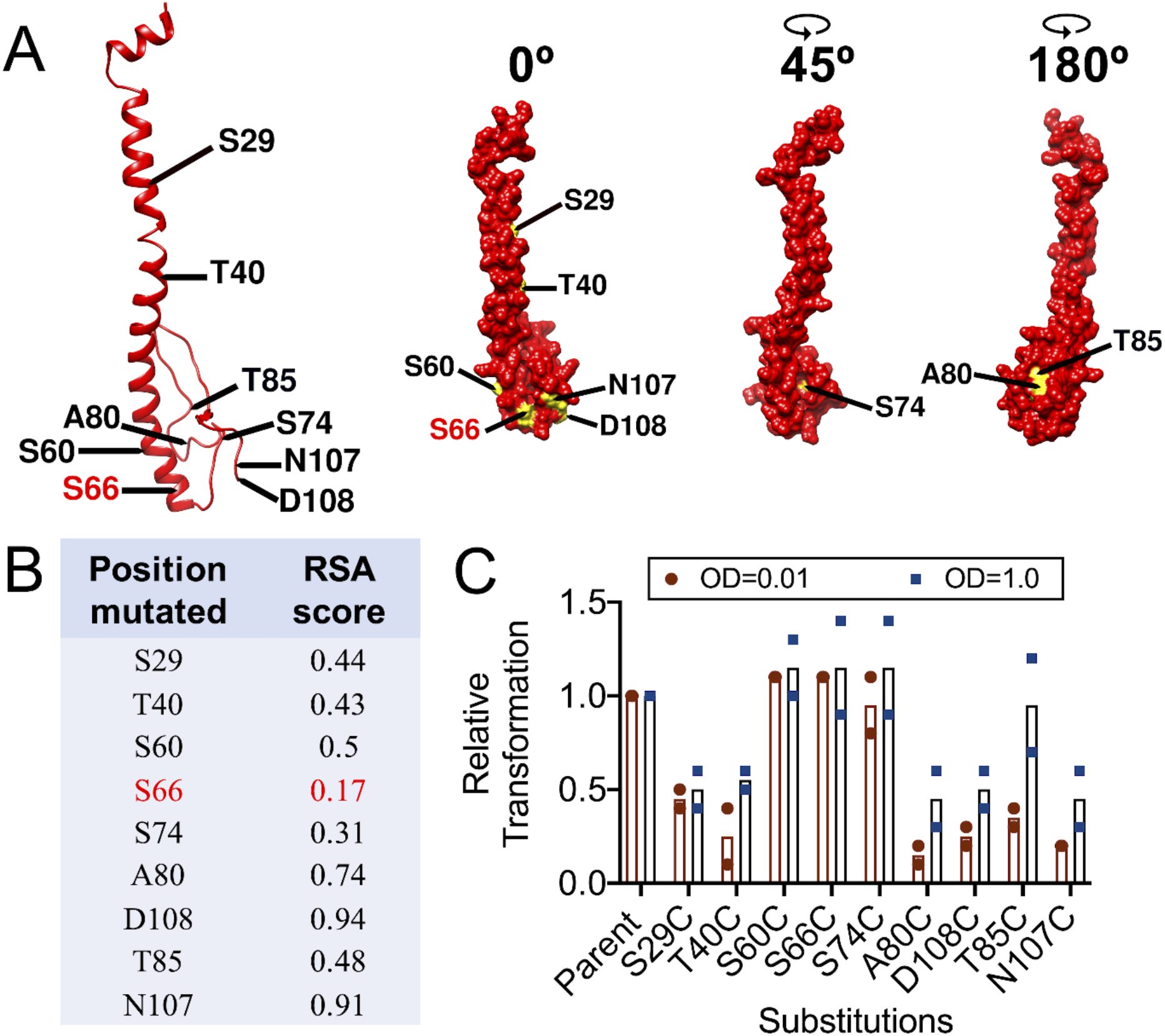
Cysteine substitution mutations at ComGC allow transformation. **A)** ComGC structure model. **Red**, cysteine replacement characterized in this paper. **B)** Relative surface accessibility (RSA) score of 9 cysteine substitution sites. **C)** Transformation efficiency of 9 *ComGC-Cys* mutants. Each strain was grown in THY to OD of 0.3, centrifuged, and resuspended in fresh THY at OD 2.0 or 0.02. Inducer was added 1:1 to each culture (OD 1.0 or 0.01), with 160 ng Nov^R^ DNA per mL. After 60 min at 37 °C, they were diluted and plated for selection. Relative transformation is calculated as a ratio of the mutant transformation efficiency to that of the parent strain. Data from each experiment (OD=0.01 and OD=1) are shown as mean and individual data points from two independent biological replicates.

### Maleimide treatment of competent cells

Type IV pili have a multistep biosynthetic provenance. Pilin subunits are synthesized ribosomally, processed by proteolysis and inserted in the cell membrane to create a pool of subunit monomers. The pilus itself is assembled from this pool of processed subunits during extrusion, while retraction reverses assembly, establishing an equilibrium between the pool of subunits in the membrane and subunits transiently incorporated in extended pili. Since the *comG* operon is expressed only in competent cells [39, 61], we reasoned that if ComG competence pili are retractile, AF488-mal tagging might best be done during an early part of a competence episode, when exposed pilin thiols would be available for reaction with the cell-impermeable maleimide reagent. After removal of unreacted dye, the accumulated AF488-tagged membrane pilin pool might continue to support further cycles of extrusion and retraction, which could be directly observed by fluorescence microscopy as dynamic pili.

Visualization of competent pneumococci labeled with AF488-mal according to this strategy would require satisfying several mutually incompatible conditions. For example, the optimal medium (chemically defined medium, CDM) for competence development includes a high level of cysteine, but cysteine quenches the maleimide reagent; and practical microscopy requires a dense cell suspension, but competence is optimal during exponential growth at low cell densities [30]. Finally, pneumococcal competence is regulated in part by an internal timer that shuts down competence expression after ~30 minutes [45, 52, 75]. Fortunately, the needed conditions can be satisfied separately but serially. Although late-log-phase or stationary-phase cells are poorly responsive to CSP [30], it was recently reported that efficient competence expression can be achieved at an artificially high cell density by collecting exponentially growing cells and simply resuspending them at a high density in fresh CDM medium [42, 43]. For this study, we took advantage of the stability of the competent state at 0-4 °C [70], to establish a labeling protocol that makes changes of medium by brief centrifugations in the cold.

Whereas competence typically reaches a maximum at ~20 min after addition of CSP [45, 61], in the adopted labeling protocol (**Fig. S1A**), exposure to CSP for 10 min at 37 °C initiates competence gene expression; then 5 min at 37 °C with AF488-mal in Cys-free CDM permits maleimide-Cys reactions as competence approaches a maximum; finally, the tagged competent cells are washed free of unreacted dye in the cold and then either imaged or exposed to donor DNA for 60 min during further incubation at 37 °C to assess residual competence. Competence of a culture that had been processed through four wash steps in the cold to effect such treatment with AF488-mal dye was compared to that of a parallel culture with no added dye, and to a culture that followed the same temperature shifts while suspended in the same complete CDM medium, but without wash steps (**Fig. S1B**). Transformation of an undisturbed CDM culture with the 5-kb Nov^R^ donor amplicon was typically ~50%; while the chilling steps of our protocol may reduce competence slightly (to ~40%), the additional washing steps did not reduce this yield further (**Fig. S1B**). While we did not determine the fraction of ComGC pilins that acquire the AF488 adduct under these conditions, the results indicate that the level of thiol modification achieved is well tolerated, with little or no effect on the fraction of cells that remain competent at the time of imaging. We conclude that this protocol yields an experimental population of AF488-mal-treated highly competent cells suitable for direct observation.

### Specific AF488-mal labeling of ComGC pilin

For direct observation of competent cells, resuspended AF488-mal-treated cells were deposited on a pre-warmed CDM agarose pad, inverted over a cover-glass, and warmed on the stage of a Deltavision Elite microscope for fluorescence imaging. This approach achieved bright cell fluorescence, at a level that was 2-3-fold higher in *ComGC*-Cys cells than in the identically treated parental strain lacking cysteine substitution, and that depended, for the *ComGC*-Cys mutant, on competence (CSP treatment) (**Fig. 2**). Thus, the majority of fluorescence signal from competent mutant cells is attributable specifically to the *ComGC*-Cys-AF488-mal adduct. The background fluorescence characteristic of the parental strain and the uninduced *comCG*-Cys mutant may reflect reaction of AF488-mal with native proteins presenting accessible thiols. Interestingly, this background label appears to be limited to the ‘newer’ surface of growing cells at the division plane (**Fig. 2A**), suggesting a maturation process that gradually exposes reactive peripheral thiol groups. For effective labeling of the competence pilus, the choice of maleimide conjugate and of Cys substitution position within ComGC are both critical (**Fig. S2**).

**Figure 2.**
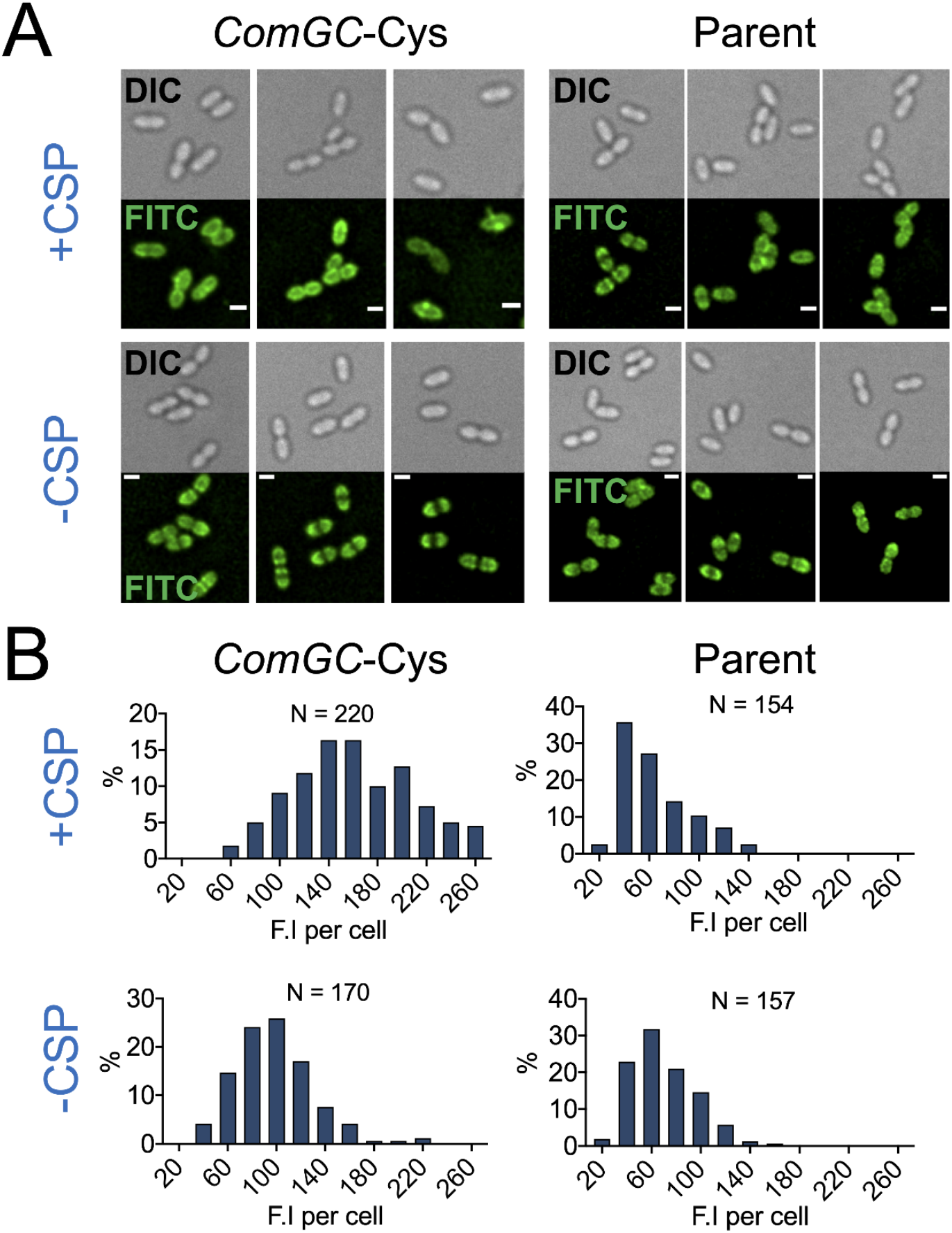
Comparison of AF488-mal fluorescence signals from competent vs. non-competent *comGC*-Cys mutant or parent strains. **A)** Representative static deconvolved images of competent (+CSP) and non-competent (−CSP) *ComGC*-Cys and parent cells labeled with AF488-mal in accord with the protocol of **Fig. S1**. Top panels, cell body imaged using differential interference contrast (DIC); bottom panels, fluorescent signal imaged using a FITC filter. Scale bars, 1 μm. **B)** Distribution of fluorescence intensity of AF488-labeled cells. The scale (%) is the percent of cells with the indicated signal among N total cells; bin width is 20.

Inspection of **Fig. 2A** shows that in the *comGC*-Cys mutant, fluorescence signal was visible in the entire cellular periphery, consistent with our expectation for equilibration of ComGC from pili into the cell membrane, both newer and older. One or two bright fluorescent foci or extended appendages also appeared specifically in many competent Cys-AF488-tagged cells (**Fig. 3 A1-A2**), but the total fluorescence signal from such cells was indistinguishable from the fluorescence signal from cells that had neither a focus nor appendage (**Fig. 3 B1-B3**).The structure of these foci was not resolved, but we speculate that they may reflect a concentration ComG pilins near the ComGB base upon which pili are assembled, or may represent short or pre-emergent pili that are not resolved by our imaging conditions. To gain some perspective on the geography of the bright foci, a cell coordinate system was adopted to map the axial positions of foci (**Fig. S3-S4** and **Fig. 3C1**). The foci were not randomly positioned, but were located predominately at the medial zone where cell wall growth occurs, and were rarely polar or subpolar (**Fig. 3C2**). We mapped the apparent bases of the extended appendages, which could be competent pili, found in competent *comGC*-Cys cells (**Fig. S5-S6**) using the same coordinate system as used above for foci. Remarkably, the apparent appendage base mapped, like foci, predominantly to the medial growth zone (**Fig. 3C3**), although polar appendages were also occasionally apparent.

**Figure 3.**
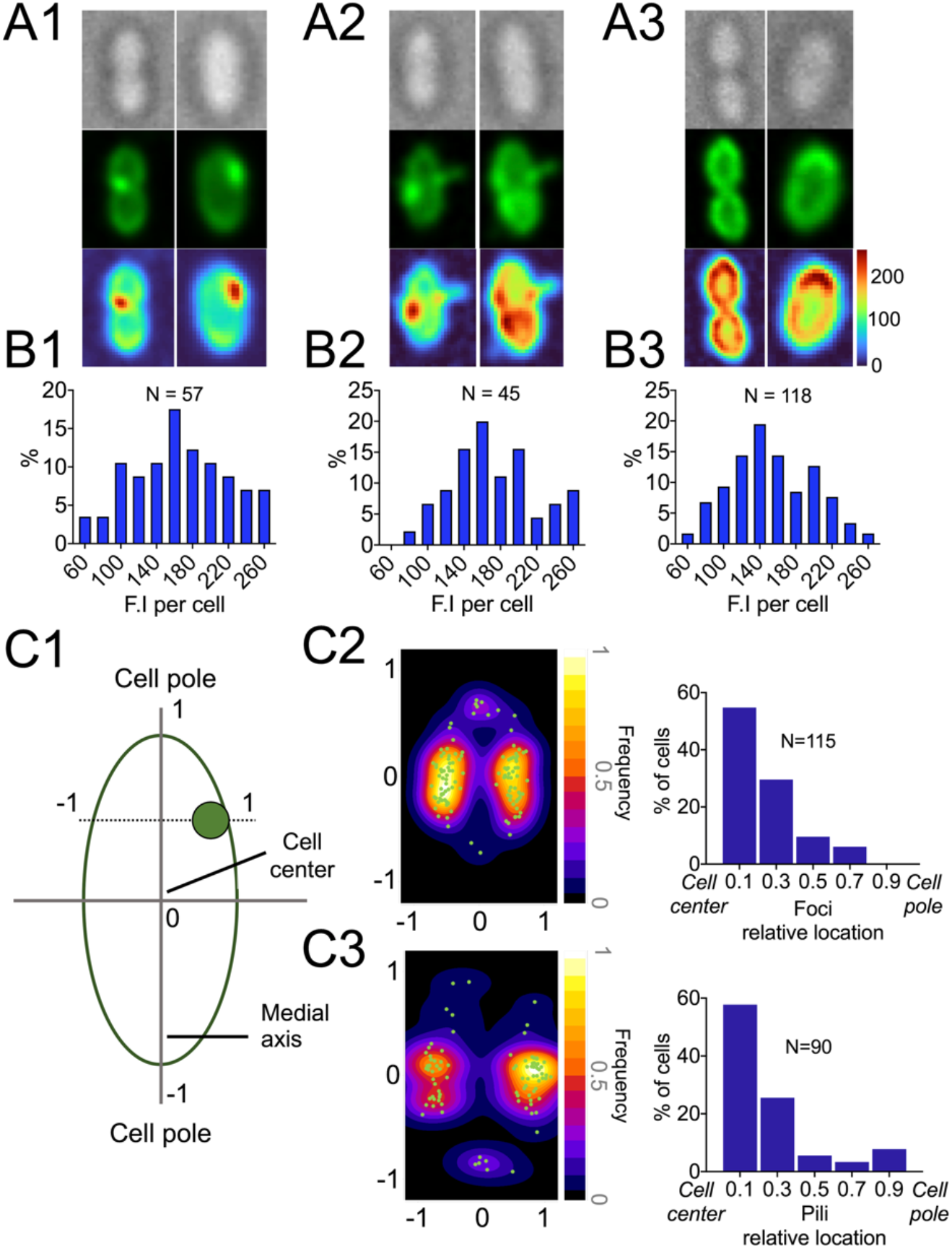
Localization of AF488-mal label in competent *ComGC*-Cys cells. Three classes of fluorescent image were distinguished: **(A1)** cells with a single bright focus; **(A2)** cells with a single extended appendage (with or without a focus); and **(A3)** cells with neither focus nor appendage. Top, DIC image; middle, deconvolved FITC image; bottom, cell fluorescence signal colormap (scale 0-255). Fluorescence intensity distribution of CSP-treated cells in each class: **(B1)** cells with a focus, **(B2)** cells with an extended appendage, or (**B3)** cells with neither. **C1**). Relative coordinates from cell center (0) to poles (−1 or 1), as used to represent focus or pilus location in C2 and C3. **C2-C3)** Localization of bright foci and extended appendages in N cells. Images that were used for the analysis are shown in **Fig. S4 and S6**. Left: heatmap of foci or extended appendages distribution; right: histogram of foci or extended appendage base location. Method of extended appendages detection is shown in **Fig. S5**. (Bin width: 0.2).

### AF488-mal-labeled protrusions extend and retract on 1-10-second timescales specifically from competent Cys-substituted cells

Noticing occasional apparently filamentous extensions in the images as described above, we searched for transient pilus-like structures associated with *comGC*-Cys competent cells, using time-lapse images of CSP-treated SAD1671 cells on the CDM agarose pads, at 2-sec intervals for a 60-sec window. Remarkably, many cells exhibited transient filamentous protrusions (**Movie S1-S4)**, which often emerged near the medial growth zone (**Fig. 3C3**) and typically disappeared, apparently by retraction, within 5-10 seconds (**Fig. 4 A1-A2**), sometimes reappearing from the same cell (**Fig. 4B**). Similar but static protrusions were also occasionally observed (**Movie S5-S7**). Pili were typically approximately 500 nm long, but occasionally over 1 micron (**Fig. 4C1**). Extrusion and retraction rates were highly variable from cell to cell and from time to time within a single extrusion episode (**Fig. 4C1**). However, retraction was regularly faster than extrusion (**Fig. 4 C1-C2, Movie S8-S15**) and a variable extrusion trajectory was typically followed by rapid retraction (**Fig. 4C2**). Fluorescent protrusions were absent from parental cells treated with AF488mal in parallel, and from non-competent SAD1671 cultures (**Fig. 4D**). Because they depended both on the *ComGC*-Cys substitution and on CSP treatment, we interpret these mobile extended structures as transiently extended *comGC*-Cys type IV pili, and conclude that the extracellular pili and pilus fragments imaged previously in fixed material from competent cultures [44, 54] are not static appendages, but represent actively mobile retractable filaments, which had been ‘frozen’ by the imaging methodology but are here revealed as such by live imaging. Since we found that the pilus base localized near midcell zone (**Fig. 3C2**), it is puzzling that the predominant midcell location of pili seen here contrasts with the [44] report that in electron micrographs of competent cells, com pili displayed no preferred location at all [44]. The discrepancy might best be resolved by applying both mapping tools in parallel to a single competent culture.

**Figure 4.**
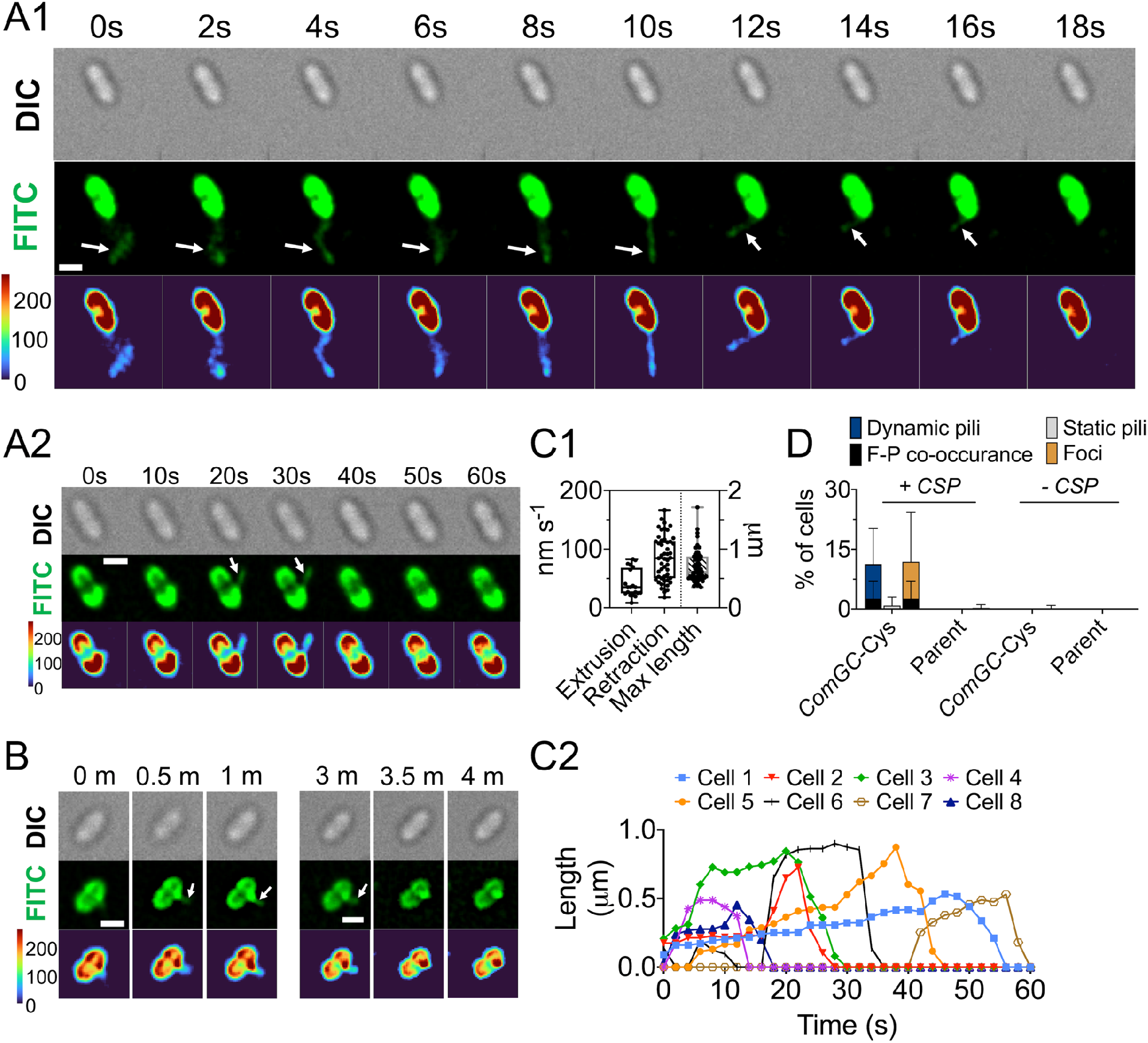
Dynamics of competence pilus activity. **(A1 and A2)** Deconvolved time-lapse imaging of *ComGC*-Cys cells with competence pilus labeled with AF488-mal. Panels: top, DIC; middle, FITC; bottom, colormap of cell fluorescence signal (scale 0-255). Arrows indicate pili. Scale bars, 1 μm. Time elapsed after capture of the first image is indicated. **B)** Persistence of *ComGC*-Cys pilus activity. Time-lapse images were taken for 1 minute with 2-sec intervals, followed by 2-minute break, and another 1-min 2-sec-interval time-lapse series of the same cell. Arrows indicate pilus. Scale bar, 1 μm. **C1)** Extrusion rate, retraction rate (nm s^-1^), and maximal length of a pilus analyzed in N cells (extrusion: N=20; retraction: N=49; max length: N=55). **C2)** Pilus dynamic kinetics of 8 representative cells. Cells that were used in the analysis are shown in **Fig. S7-S14** and **Movies S8-S15**. **D)** Quantification of time-lapse images of competent (+CSP) and non-competent (−CSP) *ComGC*-Cys and parental cells labeled with AF488-mal (N=2332 and N=510 for competent *ComGC*-Cys and parent cells, respectively; N=609 and N=629 for non-competent *ComGC*-Cys and parent cells, respectively). The percent (%) of cells indicates the fraction of cells with mobile pili, static pili, or foci among the total number of cells per frame within a 1-minute time period.

### Both transformation and retraction are blocked by neutravidin in the ComGC-Cys mutant

We reasoned that the retractable competence-specific pilus of pneumococcus might function like the type IV competence pilus of the Gram-negative pathogen, *Vibrio cholerae*, despite the long evolutionary distance between Gram-positive and Gram-negative bacteria, and that retraction of this pilus could be similarly important for DNA uptake. To test this possibility more directly, we asked whether blocking pilus retraction by attaching a bulky adduct to the *comGC*-Cys pilin would affect DNA uptake. For this purpose, a biotin-maleimide reagent (biotin-mal) was used (along with AF488-mal at a ratio of 1:5) to tag the ComGC pilins. As expected, the competition by the non-fluorescent biotin-mal reagent reduced AF488-mal labeling significantly (**Fig. 5 and Fig. S15**), but the biotin-tagged cells exhibited unaffected transformation yields and displayed mobile pili (**Fig. 5 A2 and D**), which is not surprising because biotin is a low-MW adduct that similarly does not affect the function of other T4P [25]. To attach a much bulkier adduct, the biotin-mal tagged cells were then exposed to neutravidin, a 60,000-dalton protein with high affinity for biotin [47]. The added neutravidin reduced transformation by five-fold, but had no effect whatsoever on parallel similarly treated competent parental strain cells, which lack ComGC thiols (**Fig. 5D**). Imaging of the biotin/neutravidin-treated cells confirmed that pilus retraction was strongly inhibited by the bulky adduct (**Fig. 5A3**), indicating an important role of pilus retraction in DNA uptake. Blockage of retraction was evidenced by a decrease in observable dynamic pili, an increase in static pili, and by appearance of a new class of cells that contained a large bolus of extracellular fluorescent material (**Fig. 5 A3-A4, and 5C**). This bolus contained a significantly higher relative fluorescence level than a focus or a pilus (**Fig. S16**). It may represent a tangle of pilus fibers, cross-linked by the neutravidin molecules. Indeed, because neutravidin is a tetradentate biotin-binding protein, and biotin-mal-tagged pili would carry many biotin residues, it can be expected that retraction would be blocked by formation of intra-pilus cross-links as well as by the simple steric hindrance introduced by the bulky neutravidin adducts. Because neutravidin inhibited transformation specifically of biotin-tagged cells and caused accumulation of external pili, we conclude that retraction is important for transformation, and further hypothesize that the Gram-positive type IV competence pilus transports DNA to and through the cell wall via its dynamic retraction.

**Figure 5.**
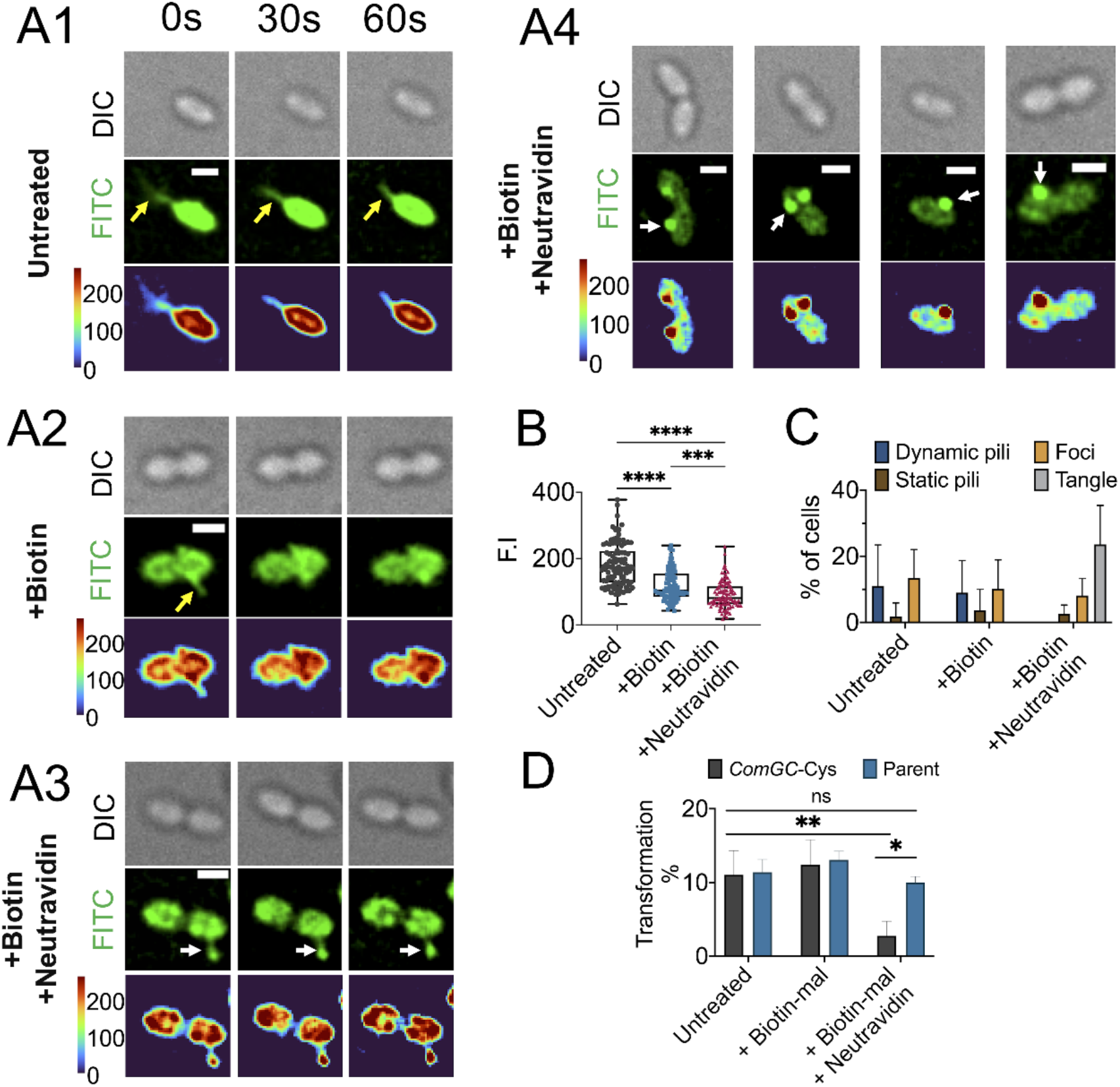
Pilus retraction is required for DNA uptake and transformation. **A1-A3)** Montage of time-lapse imaging of *ComGC*-Cys cells labeled with **A1**) AF488-mal, **A2**) biotin-mal plus AF488-mal, or **A3**) biotin-mal, AF488mal, and Neutravidin. **A4**) Representative static images of pilus tangle found in cells treated with biotin and Neutravidin. Top panels, DIC; middle panels, fluorescent images with FITC filter; bottom panels, colormap created for fluorescence signals. White arrows, static pili or tangle pili. Yellow arrows: mobile pili. Scale bar, 1 μm. **B)** Comparison of fluorescence signal among *ComGC*-Cys cells after the indicated treatments (N=100 for each condition, see supplemetal **Fig. S15 and S16**). **C**) Quantification of time-lapse images of *ComGC*-Cys competent cells labeled with AF488-mal (untreated, N=153), biotin-mal plus AF488-mal (+biotin, N=143), or biotin-mal, AF488mal, and Neutravidin (+biotin + neutravidin, N=242). The percent (%) of cells indicates the fraction of cells with dynamic pili, static pili, foci, and tangle pili among the total number of cells per frame within a 1 −minute time period. **D)** Effects of Neutravidin on transformation efficiency in *ComGC*-Cys and parent strains. Statistical analysis was done with one way ANOVA with Tukey’s post-hoc analysis (ns: non-significant, ****P<0.0001, **P<0.01).

To test this hypothesis further, we sought to visualize the interaction of pneumococcal type IV competence pili and DNA directly (**Movie S16-S17**). We used lambda phage DNA marked with the Cy5 fluorophore, which is incompatible with trans-membrane transport to the cytoplasm [6], but is readily tracked in the extracellular space. Depositing the labeled lambda DNA with competent cells on agarose pads allowed use of two-color epifluorescence to image and count both cells and DNA (**Fig. 6 A-B**). The DNA was readily visualized as individual molecules either as compact bundles or, rarely, as extended molecules. While cells were immobile on the agarose pads, the lambda DNA molecules roamed the pad, by random Brownian motion (**Movie S18-S19)**. Remarkably, DNA molecules were observed as transiently captured at the surface of 10-20% of cells. This capture occurred both with the *ComGC*-Cys mutant and with unsubstituted parent strain cells (**Fig. 6 B1-B2**), but not in either strain absent CSP treatment (**Fig. 6C**). To explore further the specificity of this capture, we mapped the subcellular location of captured DNA by mapping its apparent ‘ attachment’ site using the same approach as described in **Fig. 3** for foci and pili. Attachment predominantly at midcell (**Fig. 6D)** is consistent with a role for competence pili in facilitating the approach of nearby DNA to the surface of competent cells.

**Figure 6.**
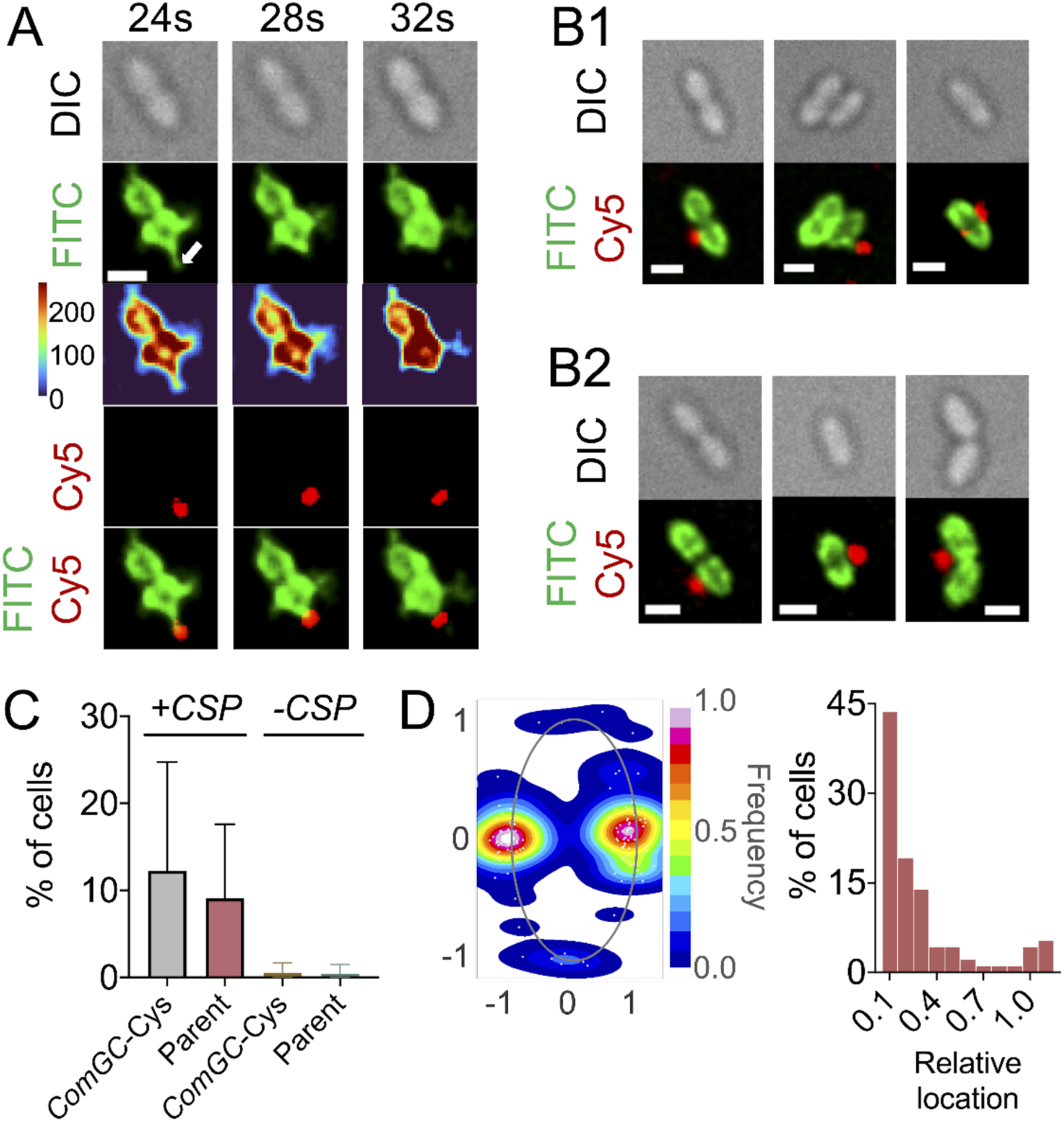
Interaction of competent *ComGC*-Cys or parent cells with Cy5-lambda DNA. A) Dynamic interaction of *ComGC*-Cys mutant with Lambda DNA (Movie S16-S17). From top to bottom: 1^st^ panel: DIC; 2^nd^ panel: fluorescent cells and pilus imaged using a FITC filter; 3rd panel: colormap created for cell and pilus fluorescence signals; 4th panel: Cy5-labeled DNA imaged using a Cy5 filter; 5th panel: merged images of Cy5 (DNA) and FITC (cells and pilus) channels. Full montage of time-lapse imaging is shown in **Fig. S17**. White arrow indicates pilus. Scale bar, 1 μm. **B)** DNA-bound competent cells of *ComGC-Cys* (**B1)** and parent strain (**B2)**. Scale bars, 1 μm. **C)** Comparison of number of cells with bound DNA for competent (+CSP) and non-competent (−CSP) *ComGC*-Cys and parent cells. DNA-bound cells were manually counted during 1-min timelapse imaging and qualified only if DNA remained close (distance from DNA fragment to cell body < 0.08 μm) to the cell body for at least 10 seconds during that time (N=2332 and N=609 for competent and non-competent *ComGC*-Cys, respectively; N=510 and N=629 for competent and non-competent parent strain, respectively). **D)** DNA binding location mapping and distribution in 113 pneumococcal cells.

## DISCUSSION

### Visualization of a mobile pneumococcal type IV competence pilus fills a gap in understanding DNA uptake in pneumococcus

Analogous to the challenge of the ‘last mile’ in domestic package delivery, the ‘last micron’ has been a persistent puzzle in describing access of competent Gram-positive bacteria to donor DNA. In the streptococci, the natural source of such DNA is understood to be lytic attack on a noncompetent cell by a nearby competent cell. Once a loop of DNA contacts a membrane-localized transport complex; endonucleolytic incision to create a double-strand break is followed by transport of one strand linearly across the membrane concomitant with reduction of the complementary strand to oligonucleotide products, by EndA. Various roles for a type IV competence pilus have been proposed in the ‘last micron’, where released DNA nears the cell periphery, passes the capsule, if present, crosses the cell wall peptidoglycan, and gains access to the periplasmic surface of the trans-membrane transport complex [64]. One possibility was that the pilus simply provides a static adhesin that accumulates DNA in the vicinity of the competent cell; others envisaged an actively retractable pilus that serves as a molecular grapnel to conduct DNA toward the cell, through the capsule, and across the cell wall either through the same opening that accommodates pilus extrusion and retraction of through an adjacent opening. To distinguish directly between static and active roles, we arranged for live-cell imaging of the pneumococcal competence pilus. The images displayed here raise serious doubt as to a purely static adhesin, while showing behavior much better suited to a grapnel-like role.

Until now, the short competence (pseudo)pilus of *B. subtilis* has offered an apparent paradigm for type IV competence pili in the Gram-positives. The ComG/PilD operons, conserved among all competent Gram-positives, comprise just eight genes. Altogether, this com pilus gene system is considerably smaller than the Gram-negative type IV competence pilus systems, where additional apparatus is devoted to pilus passage across the outer membrane [10, 35]. It would not have been surprising if the pneumococcal competence pilus acted similarly to the *B. subtilis* pseudopilus. Both long and short competence pili are known in Gram-negative species; the long Gram-negative competence pilus of *Vibrio cholerae* is retractable and brings DNA to the competent cell after binding DNA at a distance from the cell surface, via its tip. But mobility of the Gram-negative short (pseudopilus) competence pilus remains an inference, lacking direct evidence. The present observations of mobile long competence pili in *S. pneumoniae* now suggest that Gram-positive competence pili similarly occur in alternative forms. Strong conservation of the comG/PilD genes across all streptococci suggests that the mobile long com pilus of pneumococcus may be a common or universal feature of the competent state across the genus Streptococcus. It also suggests that long retractable Gram-positive competence pili may be found in multiple genera, as are their Gram-negative relatives.

It is interesting to consider whether the two pilus forms may solve different biological challenges. Elementary physical considerations suggest that a long reversibly extendable pilus with a DNA-binding tip could be especially valuable for a species, such as pneumococcus, that is often enrobed in a viscous negatively charged carbohydrate capsule that may be up to 1000 nm thick. If the capsule confronts extracellular DNA with both charge repulsion and a macromolecular impediment to diffusion, an active capsule-penetrating pilus might be doubly favorable, if it could usher DNA through not only the capsule but also through the thick peptidoglycan cell wall.

### Visualization of motility of the pneumococcal type IV competence pili leaves open many questions about the mechanism of DNA uptake

Major challenges remain for defining the path of DNA into naturally competent cells [64]. Does DNA accompany the pilus fiber across three barriers? Or follow the retracting pilus thru a residual pore? Or benefit from access to yet another, still unknown ‘pore’? Does EndA have any role in untangling potential DNA knots, or is that managed entirely by the moving pilus, prior to its passing across the cell wall? Reversible extension of the pneumococcal type IV competence pilus draws attention to the lack of an apparent retraction ATPase in all Gram-positive type IV competence operons [44]. One solution proposed is that ATPase activity of ComFA might supply the missing retraction ATPase [23]. However, pneumococcal ComFA has been isolated and biochemically characterized as a ssDNA-binding protein with ssDNA-dependent ATPase activity that interacts with itself, DprA, and ComFC, but not ComGA [22]. It would be surprising of such a small protein had a second, independent function, in pilus retraction. An attractive alternative is that the single ATPase uniformly associated with comG pilus loci acts as a reversible ATPase, much like CpaF, the single ATPase that powers motion of tad type IV pili during both extrusion and retraction in *Caulobacter crescentus* [27].

A surprising behavior of the pneumococcal competence pilus was that avidin-complexed pili regularly formed large pilus tangles consuming much of the available labeled pilin monomer. The tetravalent avidin adduct may be expected to act as an extracellular rachet that prevents pilus back-sliding during extrusion but allows continued extrusion unhindered; this could supply an ever-growing pilus of extraordinary size, attached to a single point of secretion. If tangles contain the equivalent of 2-3 microns of pilus, apparently extruded from a single site, this indicates that extrusion is not inherently limited to the typical 0.5-1 micron length, but is usually interrupted by an (unknows) length-restricting or retraction-triggering mechanism. In *Vibrio cholerae*, no similar tangles were noted on cells treated with avidin to block retraction. Because tangles of this size were not found for *V. cholerae* competence pili after blockage with avidin, we speculate that the pneumococcal com pili are more flexible that those of *V. cholerae*. Indeed, inspection of live video movies of moving pili and of electron micrographs of fixed pili shows noticeably more flexibility in the com pili from pneumococcus that for the Vibrio competence pilus [25, 44, 55].

The roles of minor pilins are unknown. Type IV competence pili are assembled as a three-start helical fiber generated by successive incorporation of pilin subunits from a membrane pool, but subunit composition and placement within the fiber are not yet well defined. Within each fiber, successive pilins are linked by a salt bridge between a conserved glutamate at the fifth position (E5) and the N-terminal amino group of the previously added pilin [20, 50, 58]. A single minor pilin lacking the conserved E5 is typical of type IV competence pili, where this subunit may be the first pilin added to the fiber and, therefore, does not require a glutamate to form a salt bridge. Mutants individually lacking each minor pneumococcal pilin are defective in transformation and pilus assembly, establishing that all four minor pilins are needed for pilus assembly [57]. Oliviera also reported observing a complex of all the minor pilins and hypothesized that a complex of minor pilins is linked to the pilus tip through the largest minor pilin, ComGG. We have not mapped DNA binding sites on the pneumococcal competence pilus; but Laurenceau *et al* [44] observed side-by-side pilus-DNA association in the EM, and a tip complex might be an alternative binding site. Finally, the role of the pilus might be different in Gram-positives, where no outer membrane pore restricts the dimensions of an incoming complex of DNA and pilus. The path of DNA through the thick peptidoglycan cell wall may accommodate the complex more easily, perhaps including the side-by-side mode of association visualized in EM micrographs by Laurenceau *et al* [44].

In several Gram-negative competent species, DNA passes through the outer membrane with participation of the periplasmic soluble protein ComEA, which acts as an entry rachet that impedes retrograde movement of DNA [31]. The rachet quickly accumulates periplasmic dsDNA, which is separately subject to nucleolytic transfer across the inner membrane. In pneumococcus and Bacillus, ComEA is not soluble, but localizes to the outer surface of the cell membrane, and is required for DNA uptake; it is not known whether it also acts as a rachet to facilitate or drive DNA movement through the cell wall into the periplasm similarly to DNA uptake in competent Gram-negative species. However, in pneumosoccus ComEA does assemble at a single focus in competent cells that recruits EndA from it normal locations throughout the membrane to a site near the site of DNA uptake [3]. The preferential mid-cell localization of ComGC foci and pili raises the question of whether they, too, are recruited by ComEA. If tDNA passes through the same wall opening as the competence pilus, colocalization of the pilus and transport proteins would potentially facilitate transfer of DNA from one to the other. However, it remains unclear whether pilus-dependent transport and trans-membrane transport are spatially or temporally coordinated events.

With the discovery that expression of CbpD, LytA, and competence bacteriocin-like peptides are linked directly to competence regulation, and with demonstration that these products increase the efficiency of transformation in mixed-cell cultures dramatically (1000-fold) [36], it became reasonable to appreciate genetic transformation as reflecting a complex coordinated system of gene transfer from one living cell to another. Specificity in the choice of victim cell is enhanced by species-specificity of CbpD’s peptidoglycan preference, the principal lysin. The rate of that transfer appears to be further enhanced by the retractable DNA binding pilus described here, either by bringing the competent cells toward the DNA mass released upon lysis of a target, or by bringing that DNA toward the competent cell, or both. Repetitious probing by a competence pilus from a single competent cell might bring multiple separate bights of the genome from a recently lysed victim to the DNA transport machine. Such a combination of DNA processing/uptake events could explain the wide scattering of recombination tracts emerging in transformants produced by 2-cell interactions [19, 43].

## Experimental procedures

### Bacterial strains and media

All pneumococcal strains used in this study were derived from the Rx1 derivative CP2137 (*hex malM511 str-1 bgl-1* Δ*comA* Δ*cps* ssbB^-^-[pEVP3]::ssbB^+^; Hex^-^ Ma^-^ ComA^-^ Cps^-^ SsbB^+^; Sm^R^ Cm^R^ low α-galactosidase background) [73]. For cell stocks, strains were grown to an optical density of 0.3 at 550 nm (OD_550_ = 0.3) in 12 mL of Todd Hewitt Broth (Becton Dickinson) with 2% yeast extract (Becton Dickinson) (THY), mixed with glycerol for a final concentration of 14%, and stored at −80 °C. CDM medium was always prepared according to Chang *et al* [4], but with addition of a 1% supplement of CAT medium [73]. When cysteine was omitted from the standard CDM formulation, the medium is denominated CDM-Cys.

### Reagents

Alexa Fluor 488 Malemide (AF488), Alexa Fluor 594 Malemide (AF594), or Alexa Fluor 647 Malemide (AF647) (Thermo Fisher Scientific) was dissolved in dimethylsulfoxide (DMSO) (Invitrogen) at 5 mg mL^-1^, 5-μL aliquoted, and stored at −20 °C. Biotin-mal (Thermo Fisher Scientific) was dissolved in DMSO at a final concentration of 5 mg mL^-1^ and sterile-filtered before 20-μL aliquots were stored at −20 °C. Neutravidin protein (Thermo Fisher Scientific) was dissolved in distilled water at of 33 mg mL^-1^, sterile-filtered, 10-μL aliquoted, and stored at −80 °C. Fluorophores were prepared and stored with limited light exposure. Lambda DNA (New England Biolabs) was labelled with Cy5 by following manufacture’s protocol (Mirus Label IT Nucleic Acid Labeling Kit Cy5), diluted to 80 μg mL^-1^ in TE buffer, aliquoted, and stored at −20 °C.

### Inducer cocktail

CSP-1 (NeoBiolab) [67] was dissolved in distilled water at 0.1 mg/ml, filtered (0.2 μm filter Sartorius), 1-mL aliquoted, and stored at −20 °C. Bovine serum albumin (BSA) (Gold Biotechnology) was dissolved at 4% (w/v) in distilled water, sterile-filtered, and stored at 4 °C. CaCl2 solid was dissolved in distilled water at 0.1 mol L^-1^, autoclaved, and stored at room temperature. An inducer cocktail containing 4 μg mL^-1^ CSP-1, 0.008% BSA, and 1 mM CaCl2 in CDM was prepared fresh before each experiment.

### Precompetent cells

Frozen stocks were diluted [1:40] in 12 mL of either THY or sterile-filtered CDM and maintained at 37°C until reaching an OD550 of 0.3. The cultures were then chilled on ice for 15 minutes, centrifuged at 7000 RPM for 7 min at 4 °C (Eppendorf model 5804R). The supernatant was discarded, and the cell pellets were resuspended in cold THY or CDM at the indicated OD, and maintained on ice until needed.

### *Labeling of comGC-Cys pilins. ComGC*-Cys cells were labeled according to Fig. S1A

Frozen cell stocks were diluted 1:40 for growth in 12 mL CDM, centrifuged, and resuspended in fresh CDM for an OD of 2. A 200-μL portion was then mixed 1:1 with inducer and incubated for 10 minutes at 37 °C to activate competence. The culture was chilled on ice for 2 minutes, centrifuged at 7000 RPM for 4 minutes, and resuspended in cold CDM-cys. This step was repeated twice before labeling with 50 μg mL^-1^ AF488-mal for 5 minutes at 37°C. Labeled cells were then centrifuged in the cold, washed twice in CDM, and resuspended in fresh CDM before imaging. To image cells and lambda DNA interaction, Cy5-labelled DNA was 10-fold diluted in CDM and mixed with 3 vols AF488-mal labeled cells before imaging. To obstruct pilus retraction, the indicated strains were co-labelled with 25 μg mL^-1^ of AF488-mal and 125 μg mL^-1^ of biotin-mal for 10 minutes at 37°C before washing twice with CDM. 4.2 μL of 33 mg mL^-1^ Neutravidin was added to the 100 μL washed cells at 1.32 mg mL^-1^ and incubated for 10 minutes on ice before imaging.

### Transformation assay

To assay competence of *comGC-Cys* mutant strains, each frozen stock was diluted, grown, centrifuged, and resuspended in THY for an OD of 2 or 0.02. The cell cultures were then mixed in 1:1 ratio with the inducer cocktail for activation of competence, followed by addition of 160 ng mL^-1^ Nov^R^ DNA and incubation for an hour at 37 °C. The resulting cultures were serially diluted and plated in soft agar as described [53] with the fourth layer containing 10 μg mL^-1^ Novobiocin for selection, or THY agar for total CFU. Plates were incubated at 37°C for at least 16 hours before counting colonies. Transformation efficiency is defined as the number of Nov^R^ transformants divided by the total viable CFU.

To determine <u>the effect of AF488-mal protocol</u> on competence (**Fig. S1B**), the strains were grown in CDM until an OD of 0.3, centrifuged, and resuspended in fresh CDM. Three cultures were prepared by mixing 1:1 ratio of cells and inducer and incubated for 10 minutes. Two of the cultures went through the labeling protocol, without and without AF488-mal added. The other culture was a positive control that followed the temperature sequences of the labeling protocol without any exchanges of medium or centrifugation. After the final wash with CDM in the labeling protocol, a saturating amount of Nov^R^ DNA (540 ng mL^-1^) was added to all cultures and incubated for an hour at 37°C before plating for Nov^R^ transformants.

To <u>observe the effect of Neutravidin (**Fig. 5D**)</u>, the indicated strains were grown in CDM, centrifuged, and resuspended in fresh CDM. Three cultures were prepared by mixing 1:1 ratio of cells and inducer and incubated for 10 minutes. Two of the cultures were co-labelled with 25 μg mL^-1^ of AF488-mal and 125 μg mL^-1^ of biotin-mal for 10 minutes and the other culture was labelled with only 25 μg mL^-1^ of AF488-mal for 10 minutes. After the last washing with CDM, NeutrAvidin (1.32 mg mL^-1^) was added to one of the co-labelled cultures. All cultures were maintained on ice for 10 minutes before 160 ng mL^-1^ of Nov^R^ DNA was added to each culture. After 60 min at 37 °C, cultures were further diluted in CAT and plated for Nov selection and CFU counts.

### Imaging and quantitative analysis

All imaging was done on 1% CDM agarose pads with a Deltavision Elite Deconvolution Microscope system (GE Healthcare Biosciences) with an environmental chamber maintained at 37 °C with a beaker of water to maintain humidity. Cell bodies were imaged using a differential interference contrast (DIC) module, whereas fluorescence was imaged using fluorescence microscopy on a Olympus IX71 microscope with Plan Apo 100X oil immersion objective, a filter set including FITC, Cy5, and mCherry filter, PCO Edge CMOS camera, and softWoRx imaging software. Cy5 labelled DNA was imaged using a Cy5 filter. Time-lapse images were acquired for 1-minute at 2-second intervals, with exposures of 150-200 msec at a light intensity of 5-10% for the fluorescent filters, 15 ms and 32% of exposure and light intensity, respectively, for DIC. For visualization, each raw image was deconvolved using built-in softWoRx imaging software, normalized for background, and corrected for photobleaching using ImageJ. For quantification, the 16-bit image at time 0 of the time-lapse imaging series was extracted and subtracted for background before measuring integrated fluorescence signal. Colormap maps were constructed in MATLAB using 16-bit deconvolved images (scale 0-255). To determine pilus maximal length, cells that had already begun retraction when imaging began or reached maximum length at the last frame were excluded from the analysis. For extrusion rate calculation, only cells with a full cycle of pilus extrusion and retraction during 1-min time-lapse imaging were used for the analysis. For retraction rate calculation, only pili retracted within a 1-min window were use for the calculation. Extrusion and retraction rate were calculated as the change in pilus length over time.

To map positions of the pilus base, foci, and DNA binding sites, deconvolved fluorescent cell images were used for detecting foci, pili, and DNA binding location along the cell medial axis in relative coordinates with cell center (0) and cell poles (1 and −1) using MicrobeJ maxima detection. Dipplococcus cell center re-adjustment is shown in **Fig. S3**. Cells with a pilus were first pre-processed using imageJ before maxima detection in microbeJ (**Fig. S5**). Heatmap distributions were created using MicrobeJ. Absolute coordinate values were taken for the distribution histogram.

### Statistics

Significance between multiple groups (>3) was calculated using One-way ANOVA with Tukey’s post-hoc analysis on Prism 9 software.

### Cysteine substitution mutagenesis

Relative surface accessibility (RSA) scores of all amino acids in ComGC sequence [54] were analyzed by Net-SurfP [60] to determine the substitutions position. ComGC structure model (**Fig. 1**) was generated from the protein sequence using Phyre2 [38] (see also **Protein Data Bank (PDB) 5NCA**) and processed for visualization using Chimera software [62].

Mutant constructs for Cys mutations were generated via splicing-by-overlap extension (SOE) PCR as previously described [18]. The upstream region of homology (UP arm; amplified with F1/R1 primers) was stitched to the downstream region of homology (DOWN arm; amplified with F2/R2 primers). Point mutations to generate the Cys mutations were incorporated into the R1/F2 primers. All mutant constructs were introduced into CP2137 by natural transformation. Output colonies were screened by colony mismatch amplification (MAMA) assay PCR [12] for the presence of the desired point mutations and confirmed by sequencing. See **Table 1** for a list of primers used to generate mutant constructs in this study.

**Table 1.**
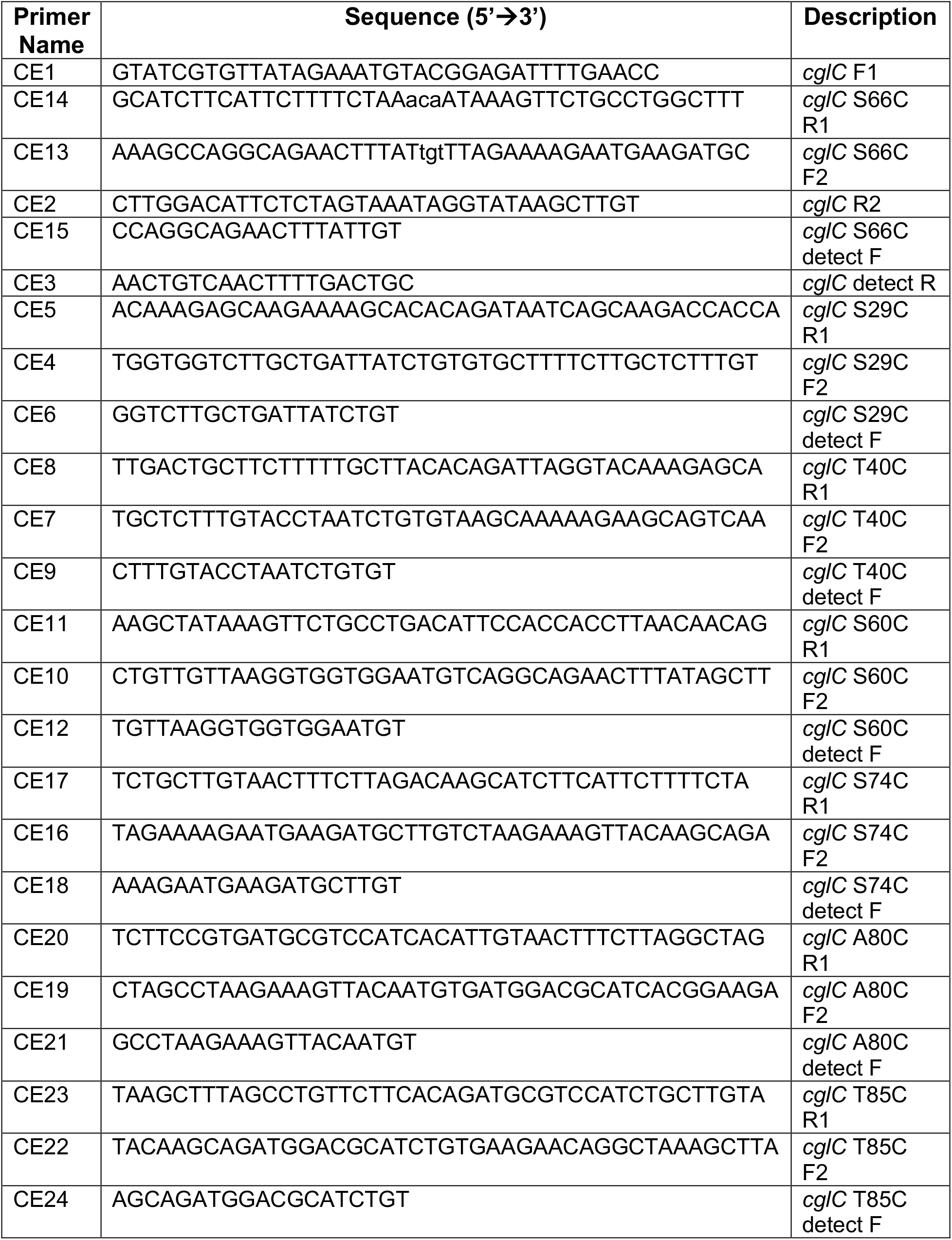
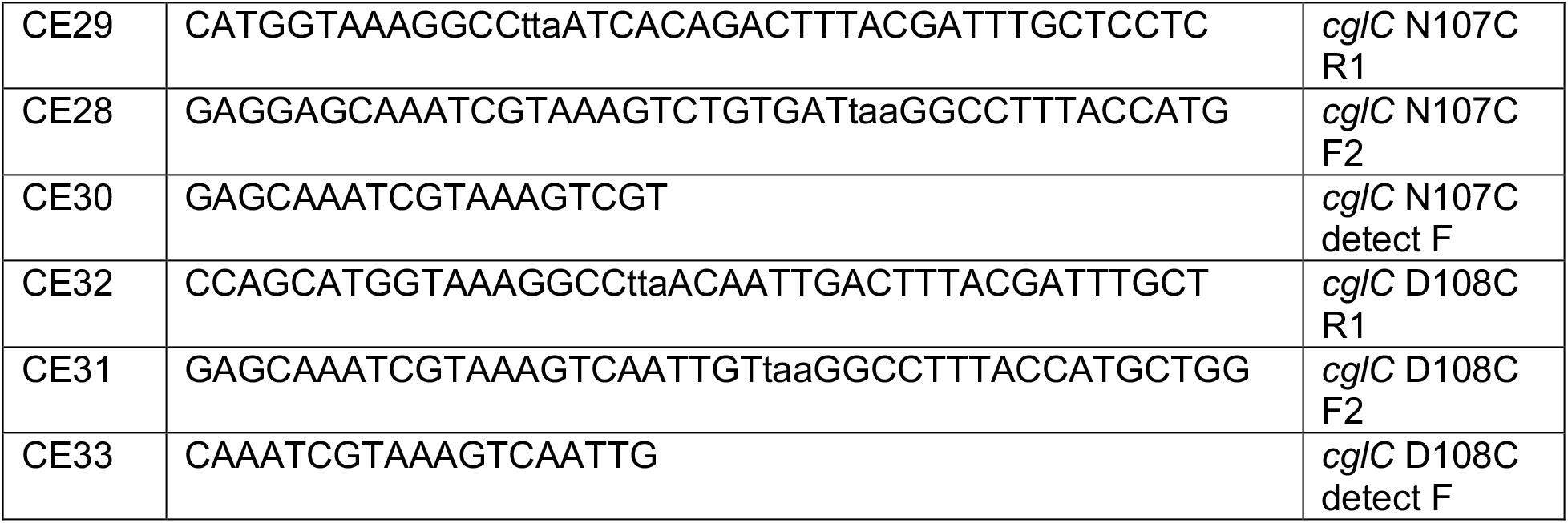
Primer oligonucleotides used in mutagenesis.

**Table 2.** Cysteine substitution mutants used in this study.

| Replacement | Strain |
| --- | --- |
| **S29C** | **SAD1668** |
| **T40C** | **SAD 1669** |
| **S60C** | **SAD 1670** |
| **<u>S66C</u>** | **SAD 1671** |
| **S74C** | **SAD 1672** |
| **A80C** | **TND 401** |
| **D108C** | **TND 402** |
| **T85C** | **TND 403** |
| **N107C** | **TND 404** |

## Acknowledgements

The authors gratefully acknowledge Xin Wang of Dr. David Stone Laboratory Department of Biological Sciences at University of Illinois at Chicago (UIC) for support of microscopy training and consultation.

