## Supplementary Information for "Type IV competence pili in *Streptococcus pneumoniae* are highly dynamic structures that retract to promote DNA uptake"

**[1]Departments of Bioengineering and [5]Biological Sciences, University of Illinois at Chicago, Chicago, IL, USA**

**[2]Lewis-Sigler Institute for Integrative Genomics and [3]Department of Molecular Biology, Princeton University, Princeton, NJ, USA**

**[4]Department of Biology, Indiana University, Bloomington, IN, USA**

**Short summary.** Competent pneumococci kill non-competent cells on contact. Retractable DNA-binding fibers in the class of type IV pili may provide a key tool for retrieving DNA segments from cell wreckage for internalization and recombination.

**Funding statement.** This work was supported by the National Institutes of Health [award R21AI133304 to DTE and award R35GM128674 to ABD] and National Science Foundation [NSF fellowship 1342962 awarded to CKE]

**Conflict of interest.** There are no conflicts to declare.

**Contributions of Authors.** AD, DM, and CE conceived and designed the study. TL performed the experiments. TL, DM, and AD analyzed the data. CE and AD contributed reagents/materials/analysis tools. DM and TL wrote the paper. AD and DE edited the paper.

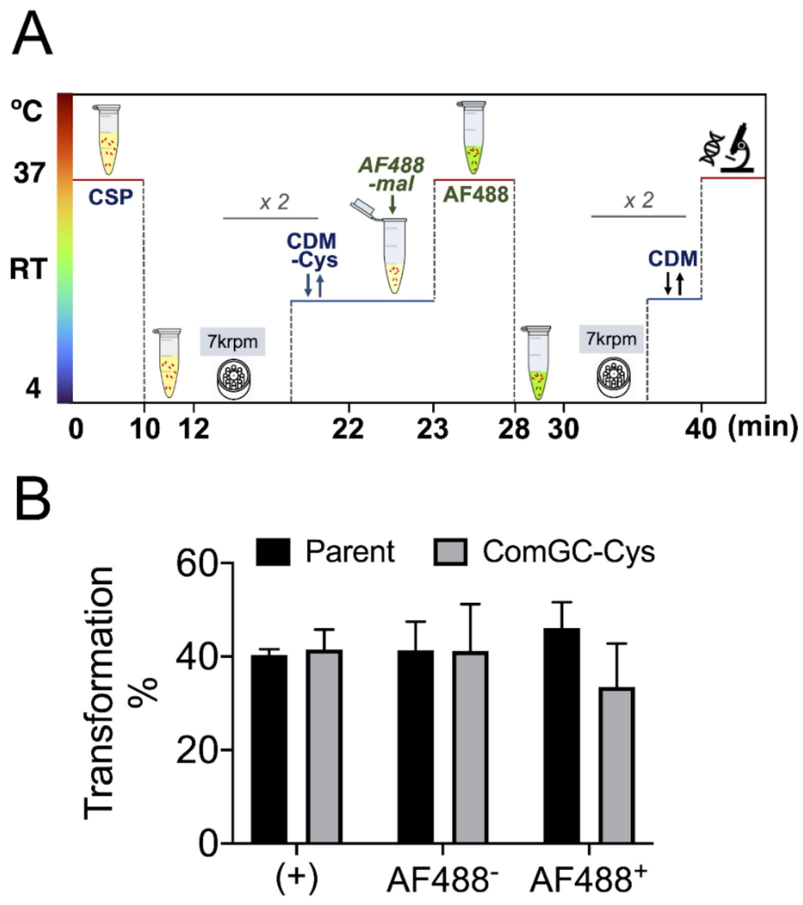


**Figure S1. Stability of competence through maleimide labeling protocol**. **A)** Labeling protocol for AF488-mal. Cells and inducer in CDM are mixed 1:1 and incubated at 37°C for 10 minutes to activate competence. Then the culture is chilled to 0°C and washed twice in the cold with fresh CDM-Cys, followed by incubation with AF488-mal in CDM-Cys at 37°C for 5 minutes. The culture was again chilled, washed twice, and resuspended in cold CDM at OD 1.0 for competence determination or imaging on a CDM agarose pad. **B)** Persistence of competence in the S66C *ComGC-Cys* mutant through washing and exchanges of media in the labeling protocol. Three aliquots of a culture of S66C (SAD1671) resuspended in CDM were prepared and labeled as positive control (+), AF488+, and AF488-. AF488+ and AF488- passed through the labeling protocol with and without AF488-mal added, respectively. The positive control (+) went through the same temperature sequence, but without washing or exchanges of media. For transformation, was each culture was supplemented with $Nov^R$ DNA (540 ng/mL), incubated for an hour at 37°C, diluted, and plated for Nov selection. The experiment was replicated thrice (N=3), and statistical comparisons were made by one-way ANOVA in Prism 9 with Tukey's post-hoc analysis. Error bars indicate standard deviation among replicates.

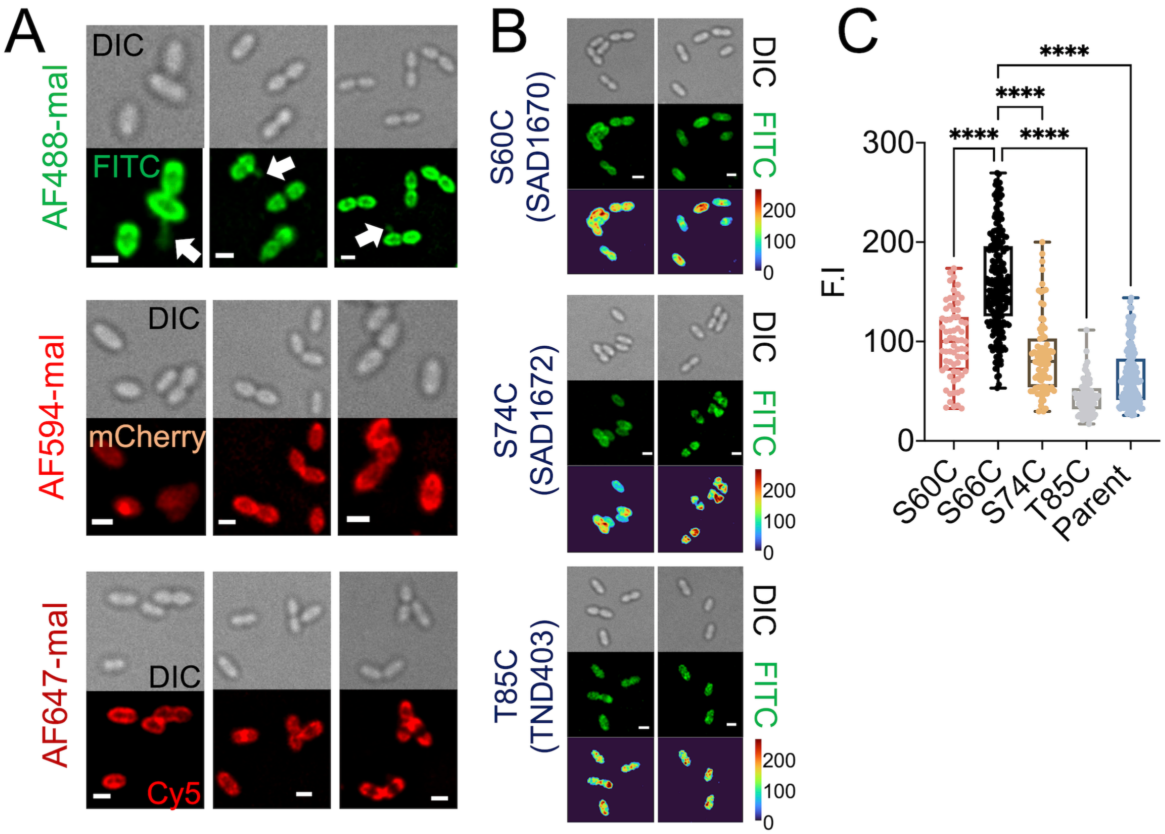


**Figure S2. Comparison of fluorescence imaging of different *comGC*-Cys mutants using various maleimide conjugates. A)** Representative static images of competent *ComGC*-Cys (SAD1671, S66C) labeled with AF488-mal, AF594-mal, or AF647-mal according to the protocol shown in **Fig. S1**. White arrows indicate pili. Scale bars, 1 µm. **B)** Imaging of different *ComGC*-Cys mutants labeled with AF488-mal. Top: S60C; middle: S74C; and bottom: T85C. Top panels: cell body imaged using DIC; middle panels: fluorescent signal imaged using a FITC filter; bottom panels: heatmap created for fluorescence signals (scale 0-255). Scale bars, 1 µm. Imaging was done on 1% agarose CDM pads in a 37ºC imaging chamber. Images were acquired and deconvolved using softWorx software of the Deltavision imaging system. The cell body was imaged with 32% light intensity and 15ms exposure of DIC, while fluorescent cells and pili were imaged with 5% light intensity and 150ms with a FITC filter (AF488), an mCherry filter (AF594), or a Cy5 filter (AF647). **C)** Comparison of fluorescence signals of different *comGC*-Cys mutants labeled with AF488-mal. Quantitative analysis used raw 16-bit TIFF files extracted from time point 0 in time-lapse imaging series and using ImageJ. The scale (%) is the percent of cells with the indicated signal among N total cells ($N_{S60C}$=69, $N_{S66C}$=220, $N_{S74C}$=85, $S_{T85C}$=78, $N_{parent}$=154).

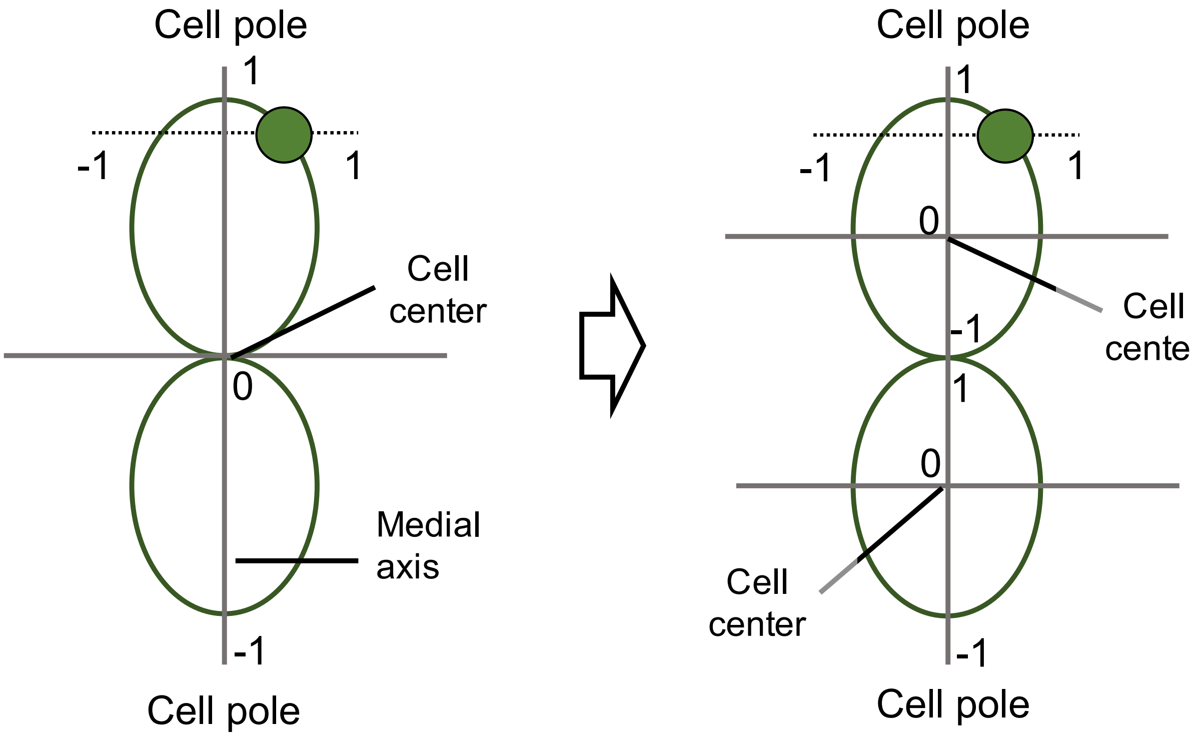


**Figure S3. Dipplococcal cell center re-adjustment using microbeJ.** Dipplococcus cell images were imported to microbeJ and segmented using microbeJ manual editing tool for cell center correction. **(Left)** Dipplococcal cell coordinates detected automatically by microbeJ before adjustment. **(Right)** Dipplococcal cells coordinates after adjustment.

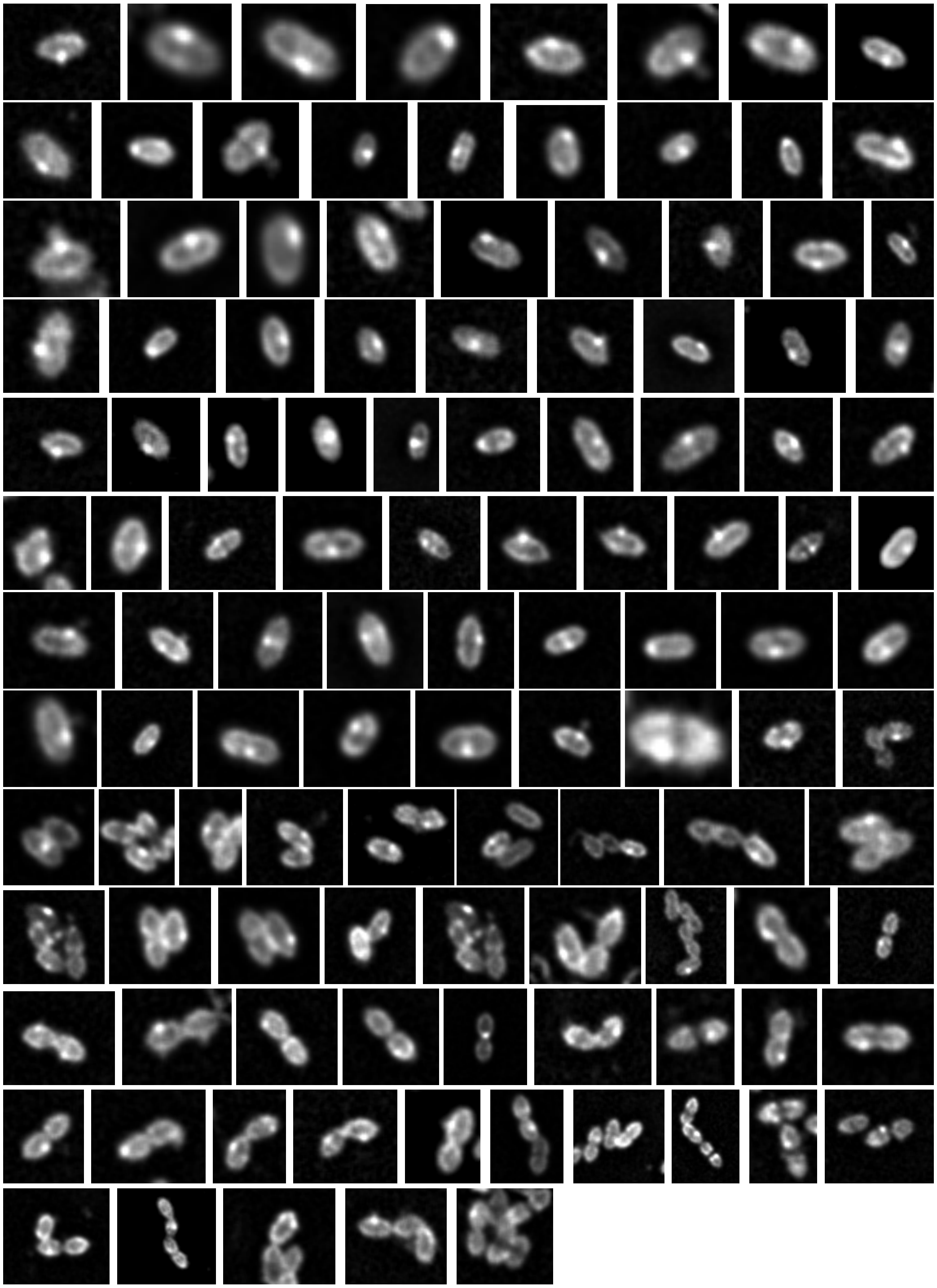

**Figure S4. Fluorescent cell images of 115 AF488-labeled competent cells that were used for foci localization.** Cells were labeled with AF488-mal in accord with the protocol of **Fig. S1**. Imaging was done on 1% agarose CDM pads in a 37ºC imaging chamber. Images were deconvolved using softWoRx software of Deltavision imaging system.

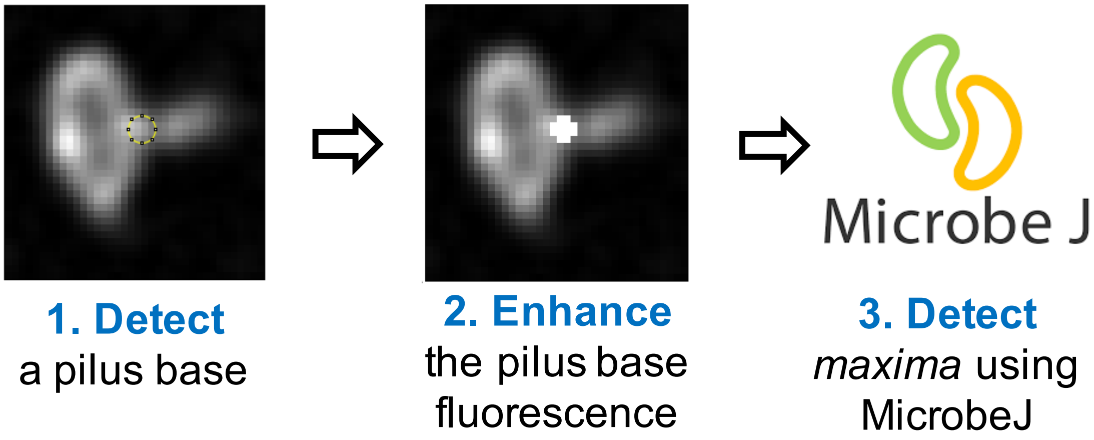


**Figure S5. Method for detecting pilus base in AF488-labeled competent *comGC*-Cys cells.** Pili-base detection was accomplished using imageJ and microbeJ. Static deconvolved images of cells with a pilus were used for the analysis. Pili base fluorescence signal was manually enhanced to saturation for maxima detection function in microbeJ.

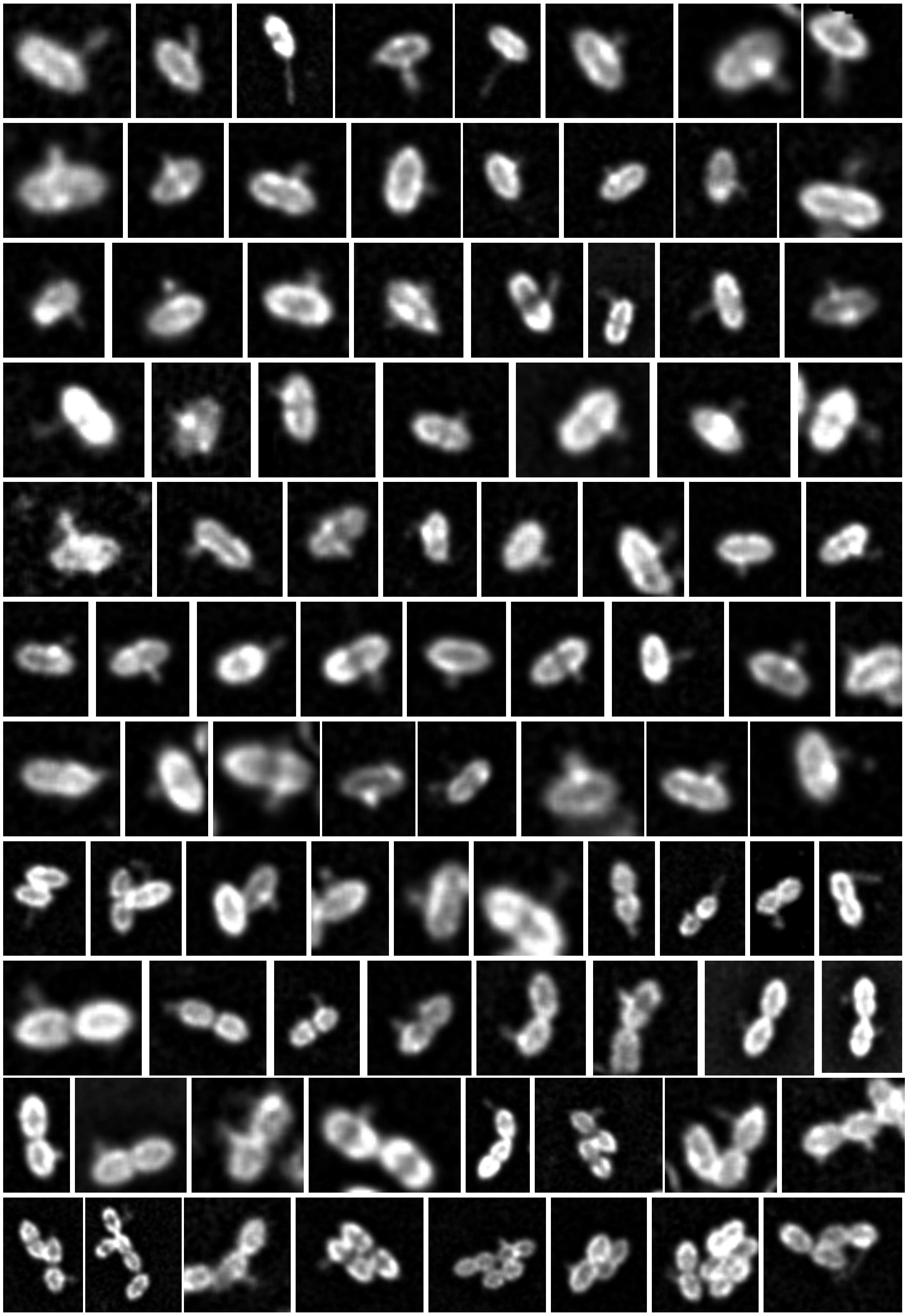

**Figure S6. Fluorescent cell images of 90 AF488-labeled competent cells that were used for pili base location detection.** Cells were labeled with AF488-mal in accord with the protocol of **Fig. S1**. Imaging was done on 1% agarose CDM pads in a 37ºC imaging chamber. Images were deconvolved using softWoRx software of Deltavision imaging system.

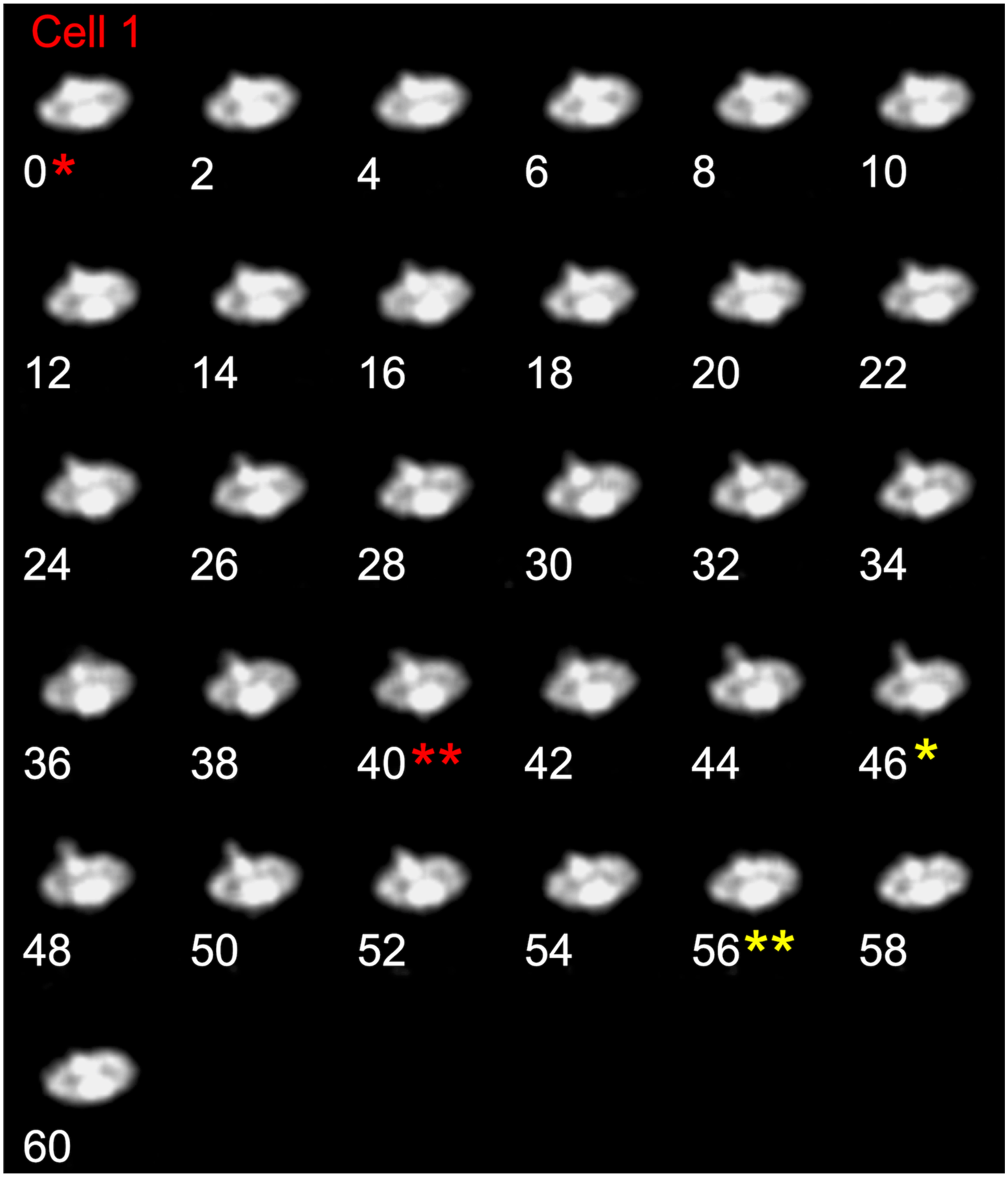


**Figure S7A. Montage of time-lapse images of the *ComGC*-Cys cell that was used in Fig. 4 C2 (Cell 1).** Cell was labeled in accord to **Fig. S1**. Time-lapse images were acquired for 1-minute at 2-second intervals, with exposures of 150 ms at a light intensity of 5% for the fluorescent filters, and 15 ms and 32% of exposure and light intensity for DIC. Red * indicates extrusion starting time and red ** indicates extrusion end time. Yellow * indicates retraction starting time and ** indicates retraction end time.

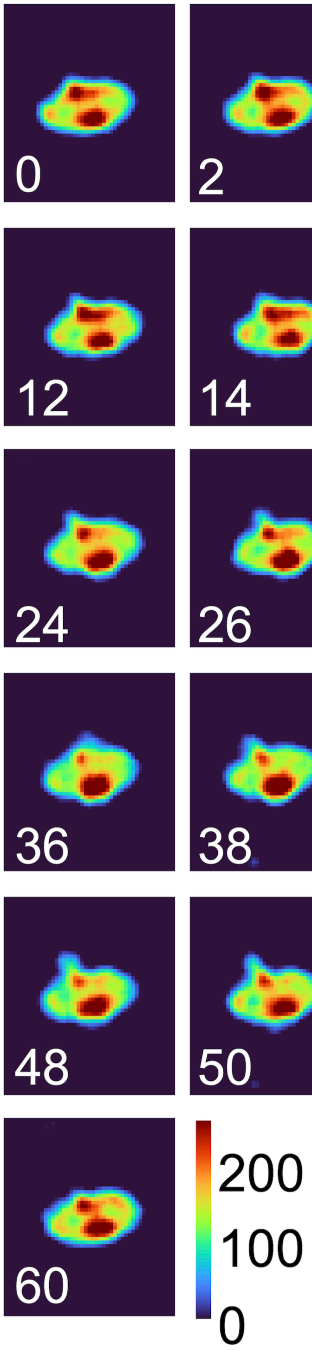


**Figure S7B. Colormap montage of time-lapse images of the *ComGC*-Cys cell that was used in Fig. 4 C2 (Cell 1).** Colormap maps were constructed in MATLAB using 16-bit deconvolved images (scale 0-255).

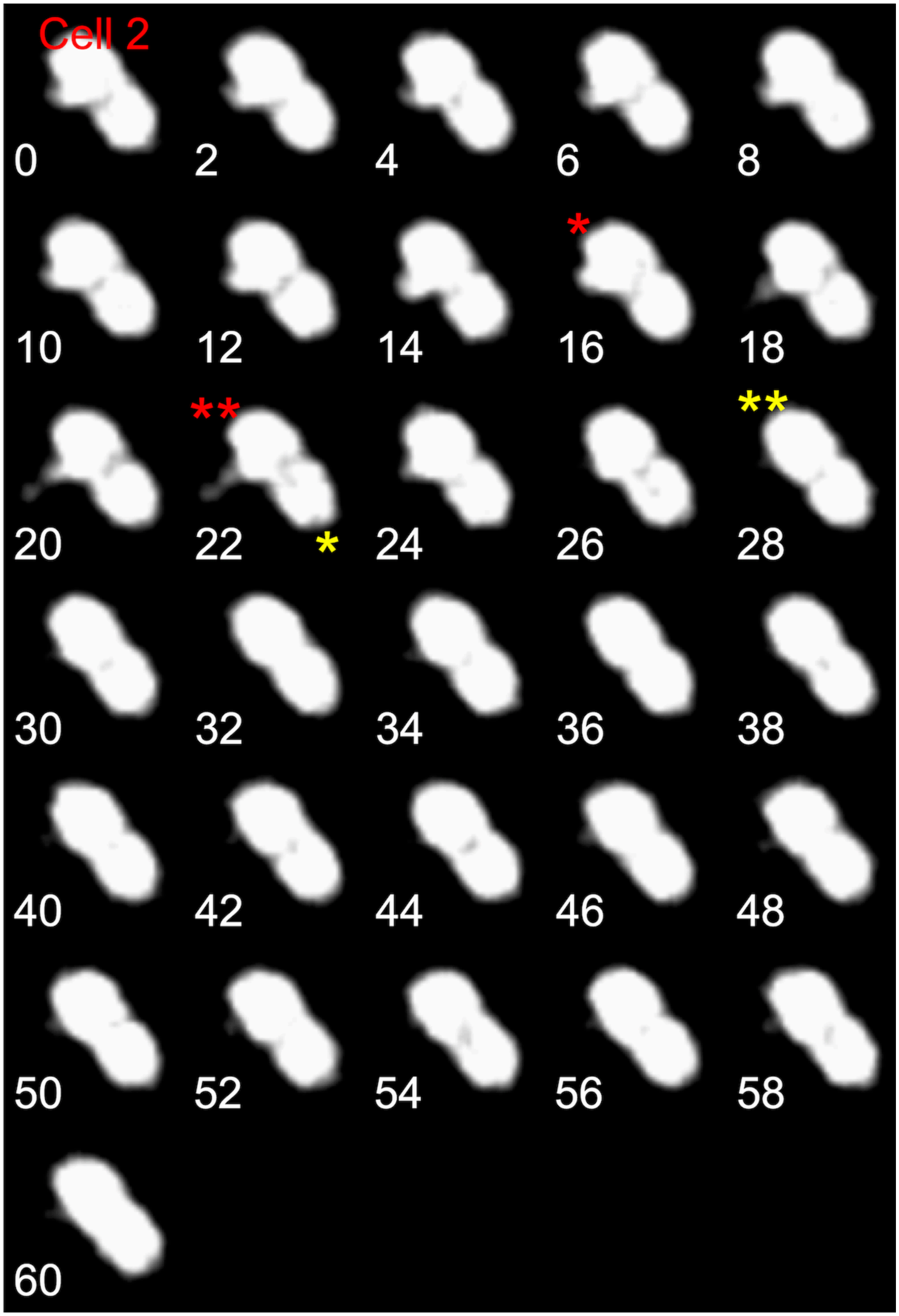


**Figure S8A. Montage of  time-lapse images of the *ComGC*-Cys cell that was used in Fig. 4 C2 (Cell 2).** Cell were labeled in accord to **Fig. S1.** Time-lapse images were acquired for 1-minute at 2-second intervals, with exposures of 150 ms at a light intensity of 5% for the fluorescent filters, and 15 ms and 32% of exposure and light intensity for DIC. Red * indicates extrusion starting time and red ** indicates extrusion end time. Yellow * indicates retraction starting time and ** indicates for retraction end time.

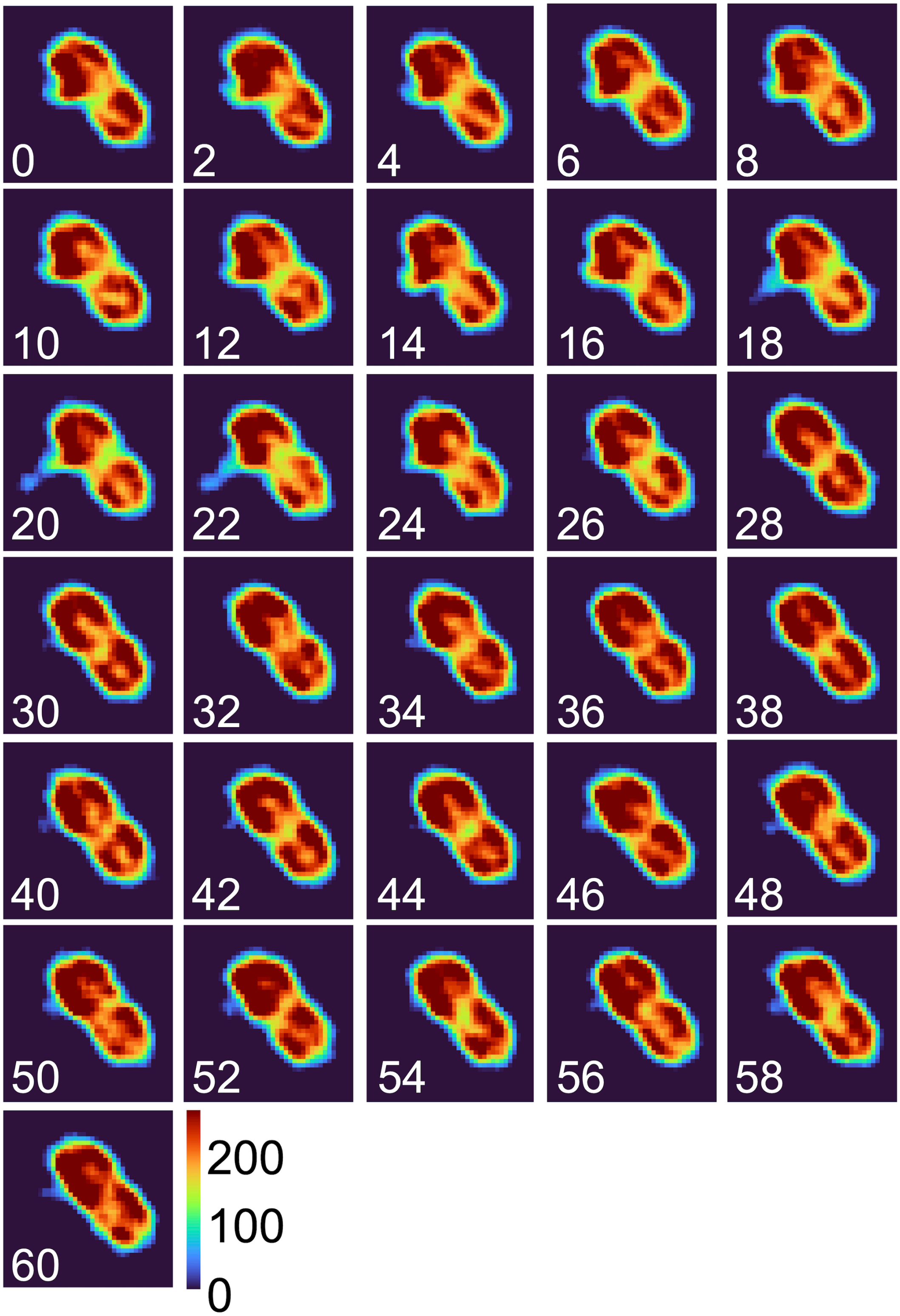


**Figure S8B. Colormap montage of time-lapse images of the *ComGC*-Cys cell that was used in Fig. 4 C2 (Cell 2).** Colormap maps were constructed in MATLAB using 16-bit deconvolved images (scale 0-255).

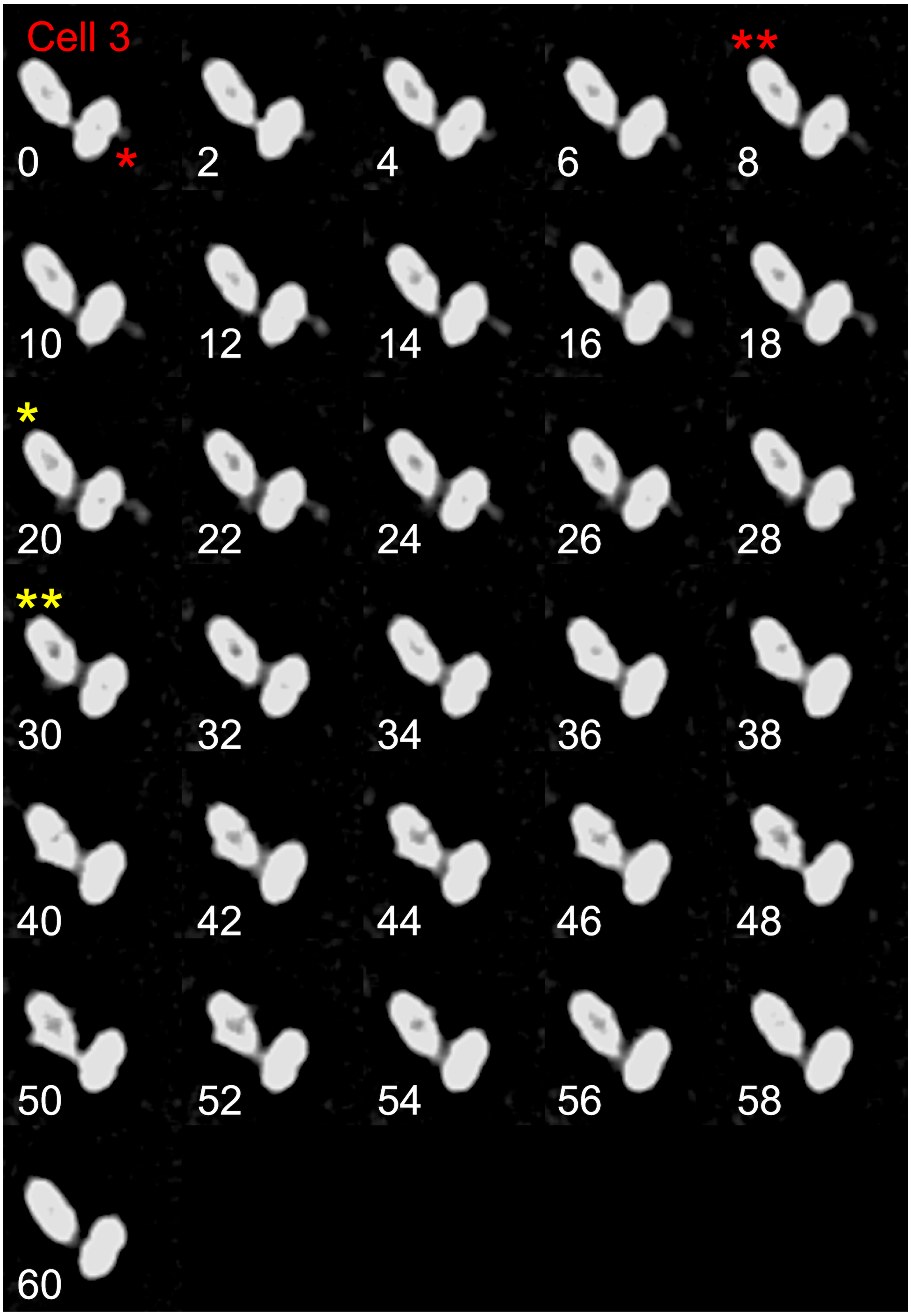


**Figure S9A. Montage of time-lapse images of the *ComGC*-Cys cell that was used in Fig. 4 C2 (Cell 3).** Cell were labeled in accord to **Fig. S1**. Time-lapse images were acquired for 1-minute at 2-second intervals, with exposures of 150 ms at a light intensity of 5% for the fluorescent filters, and 15 ms and 32% of exposure and light intensity for DIC. Red * indicates extrusion starting time and red ** indicates extrusion end time. Yellow * indicates retraction starting time and ** indicates for retraction end time.

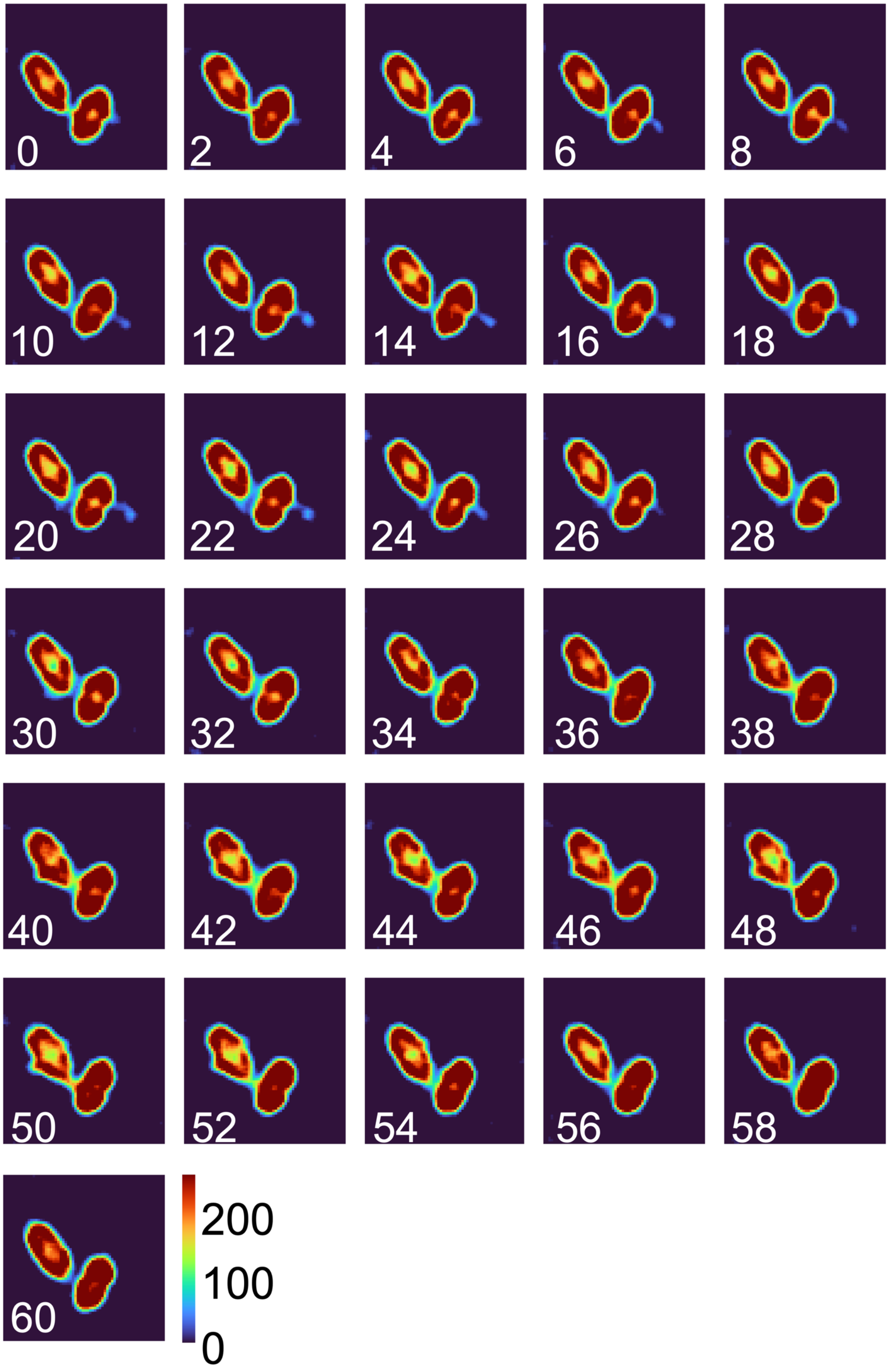


**Figure S9B. Colormap montage of time-lapse images of the *ComGC*-Cys cell that was used in Fig. 4 C2 (Cell 3).** Colormap maps were constructed in MATLAB using 16-bit deconvolved images (scale 0-255).

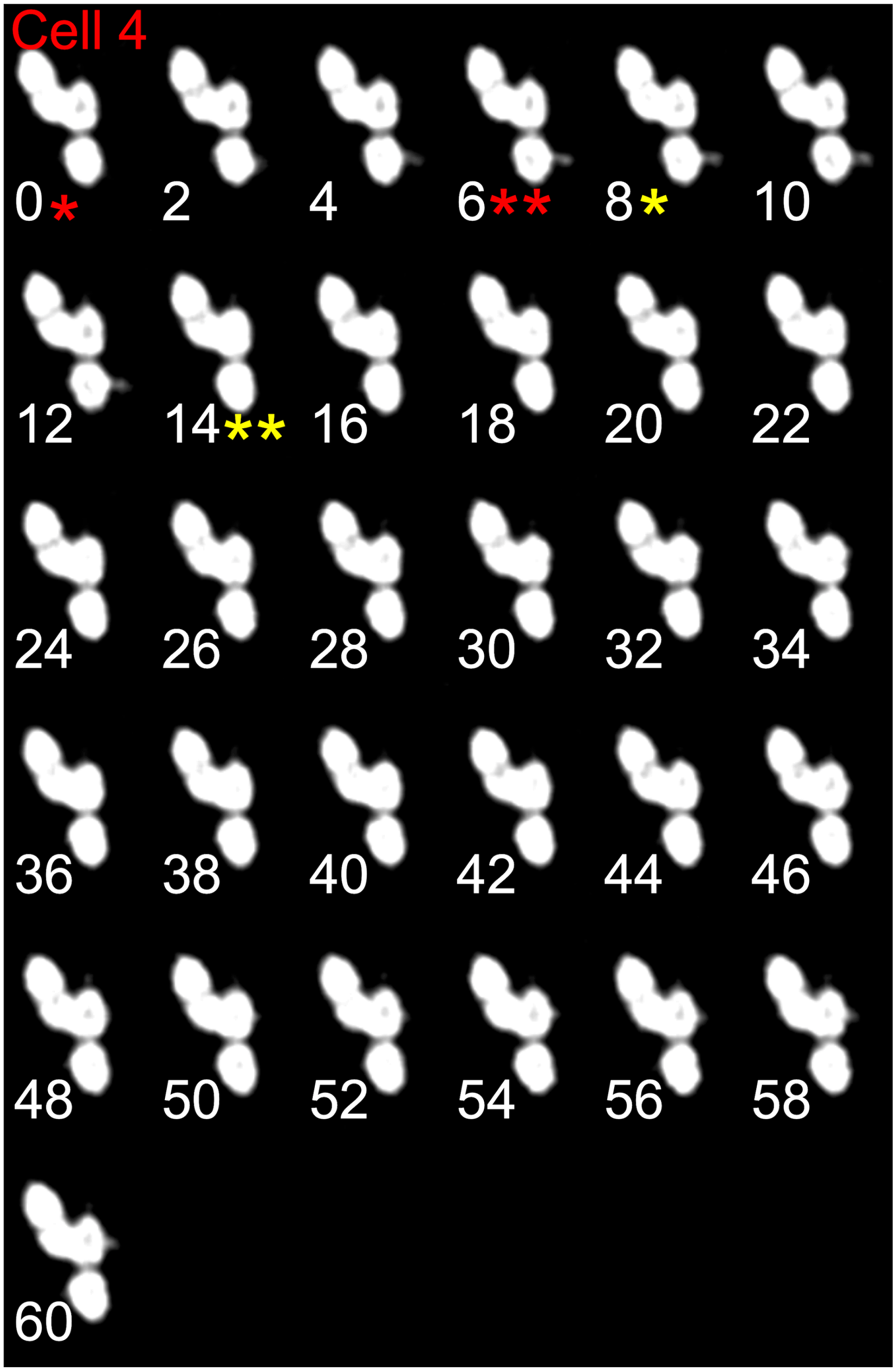


**Figure S10A. Montages of time-lapse imaging of the *ComGC*-Cys cell that was used in Fig. 4 C2 (Cell 4).** Cell were labeled in accord to **Fig. S1**. Time-lapse images were acquired for 1-minute at 2-second intervals, with exposures of 150 ms at a light intensity of 5% for the fluorescent filters, and 15 ms and 32% of exposure and light intensity for DIC. Red * indicates extrusion starting time and red ** indicates extrusion end time. Yellow * indicates retraction starting time and ** indicates for retraction end time.

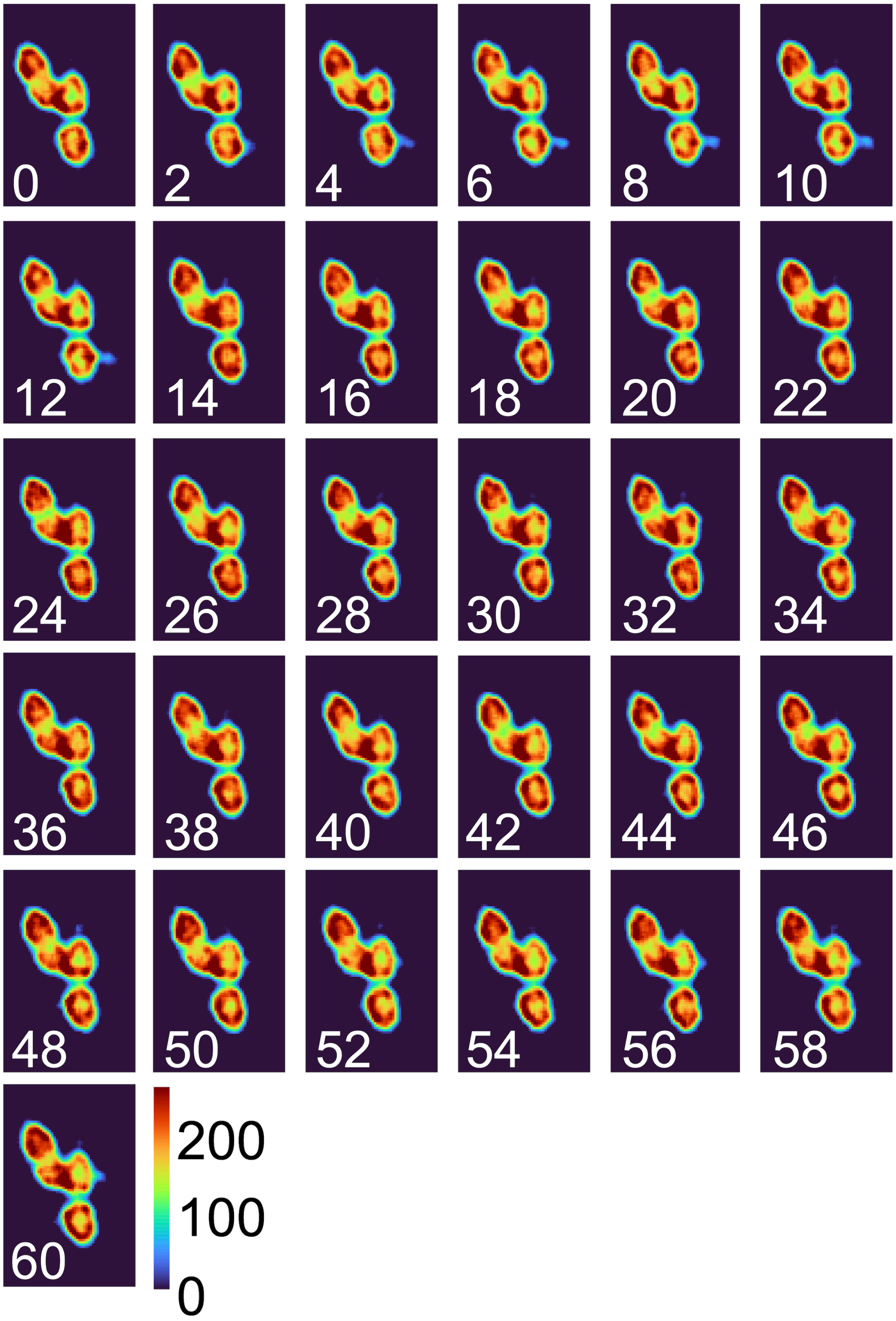


**Figure S10B. Colormap montage of time-lapse images of the *ComGC*-Cys cell that was used in Fig. 4 C2 (Cell 4).** Colormap maps were constructed in MATLAB using 16-bit deconvolved images (scale 0-255).

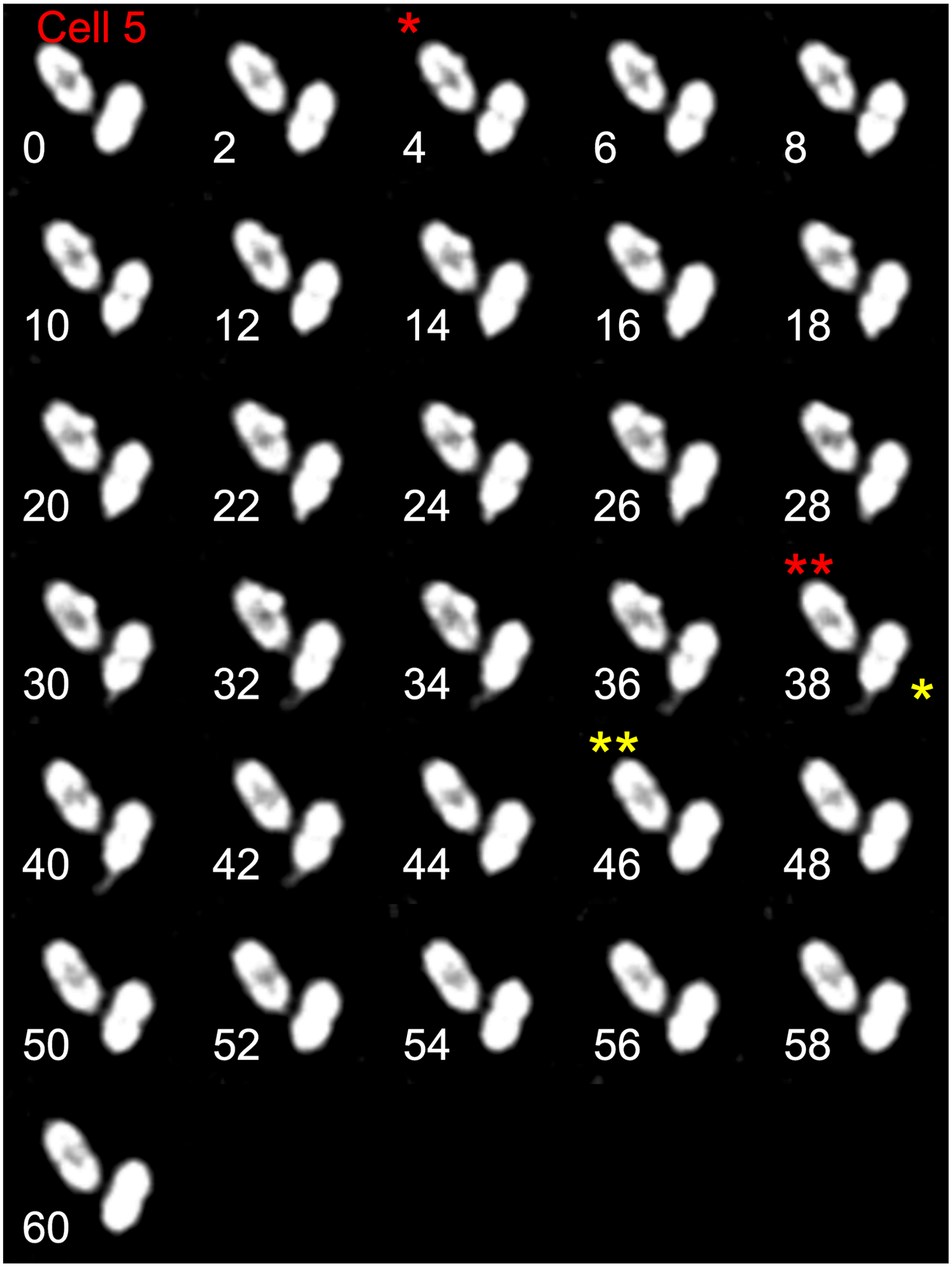


**Figure S11A. Montages of time-lapse imaging of the *ComGC*-Cys cell that was used in Fig. 4 C2 (Cell 5).** Cell were labeled in accord to **Fig. S1**. Time-lapse images were acquired for 1-minute at 2-second intervals, with exposures of 150 ms at a light intensity of 5% for the fluorescent filters, and 15 ms and 32% of exposure and light intensity for DIC. Red * indicates extrusion starting time and red ** indicates extrusion end time. Yellow * indicates retraction starting time and ** indicates for retraction end time.

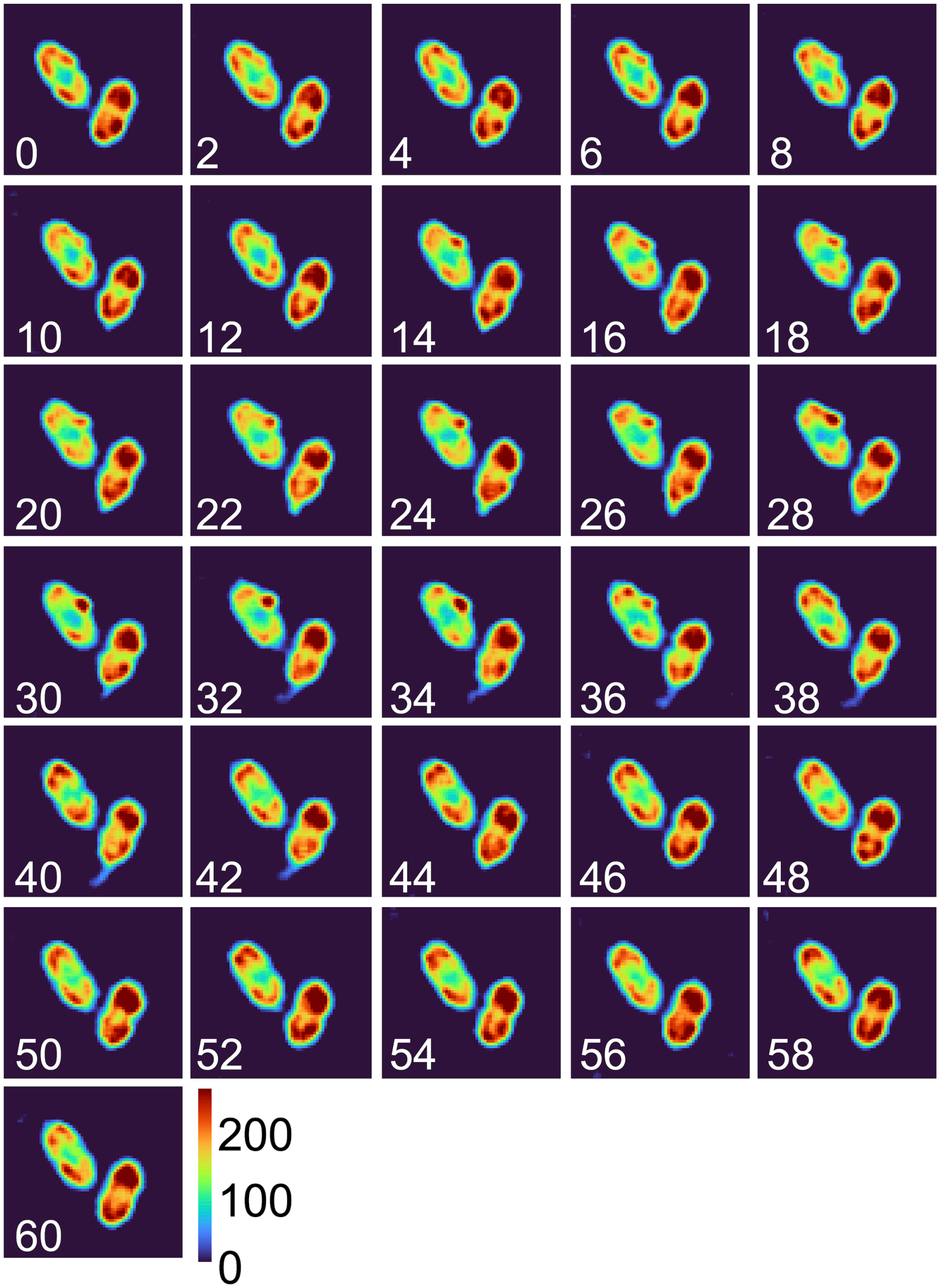


**Figure S11B. Colormap montage of time-lapse images of the *ComGC*-Cys cell that was used in Fig. 4 C2 (Cell 5).** Colormap maps were constructed in MATLAB using 16-bit deconvolved images (scale 0-255).

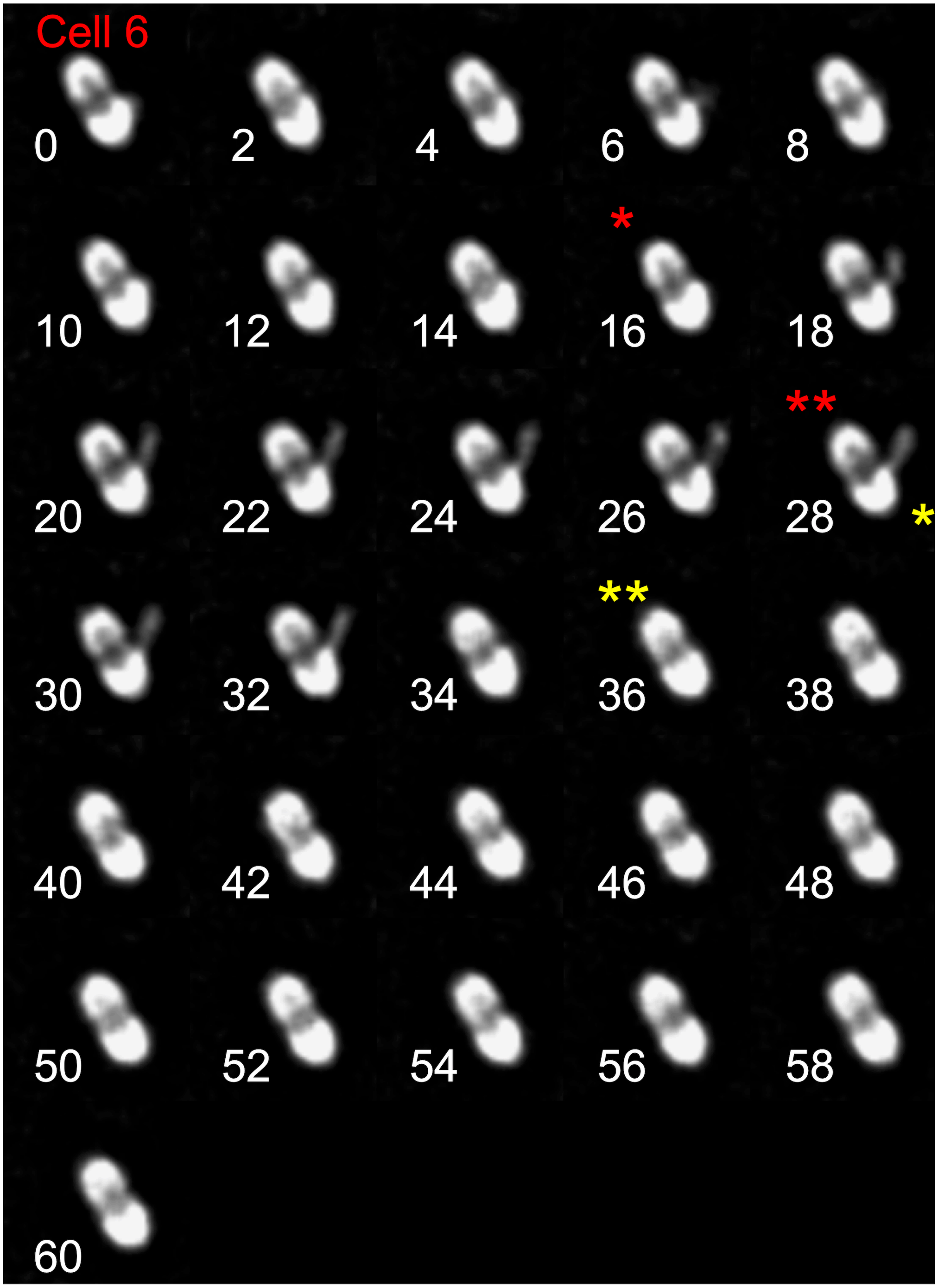


**Figure S12A. Montages of time-lapse imaging of the *ComGC*-Cys cell that was used in Fig. 4 C2 (Cell 6).** Cell were labeled in accord to **Fig. S1**. Time-lapse images were acquired for 1-minute at 2-second intervals, with exposures of 150 ms at a light intensity of 5% for the fluorescent filters, and 15 ms and 32% of exposure and light intensity for DIC. Red * indicates extrusion starting time and red ** indicates extrusion end time. Yellow * indicates retraction starting time and ** indicates for retraction end time.

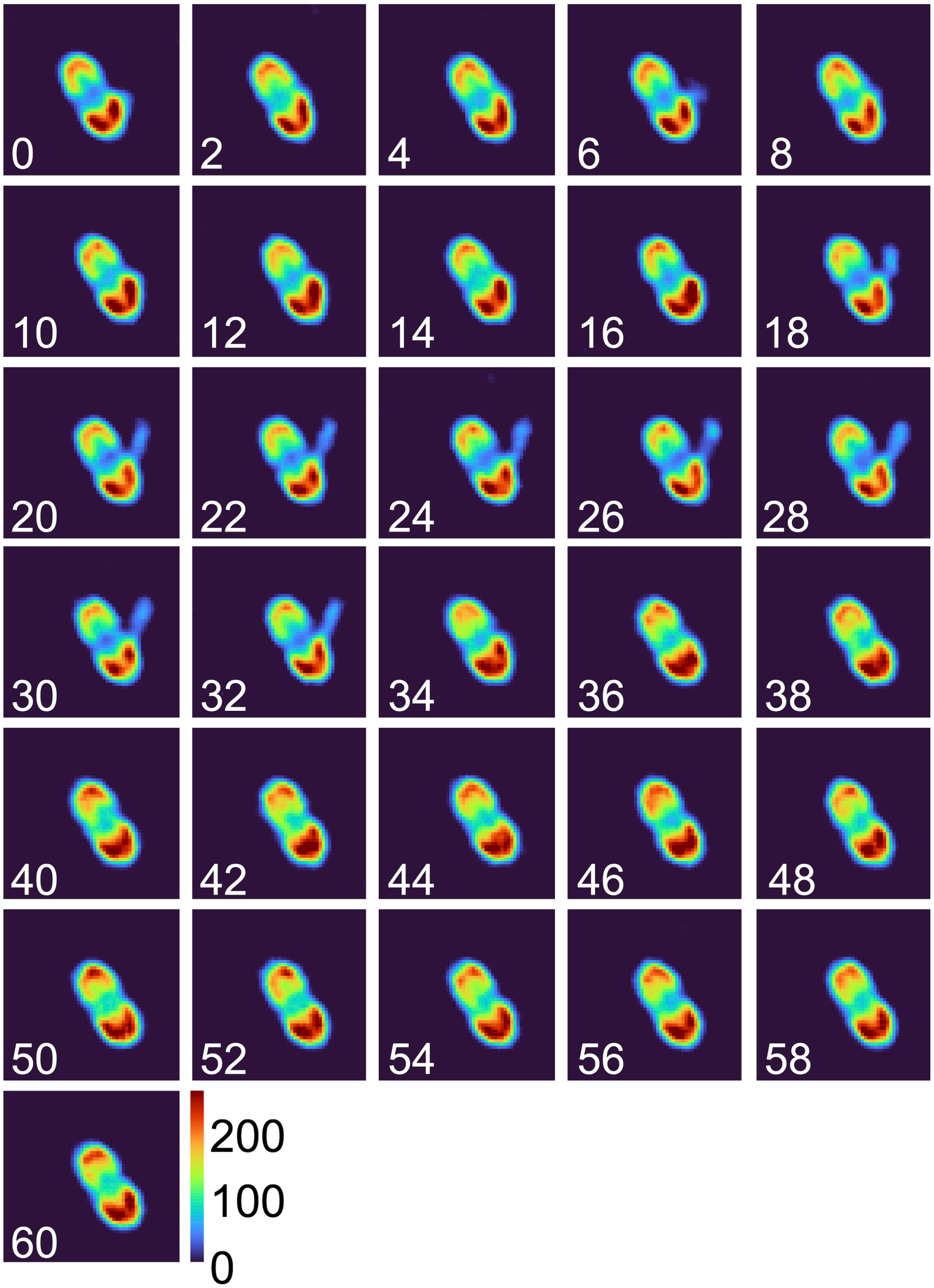


**Figure S12B. Colormap montage of time-lapse images of the *ComGC*-Cys cell that was used in Fig. 4 C2 (Cell 6).** Colormap maps were constructed in MATLAB using 16-bit deconvolved images (scale 0-255).

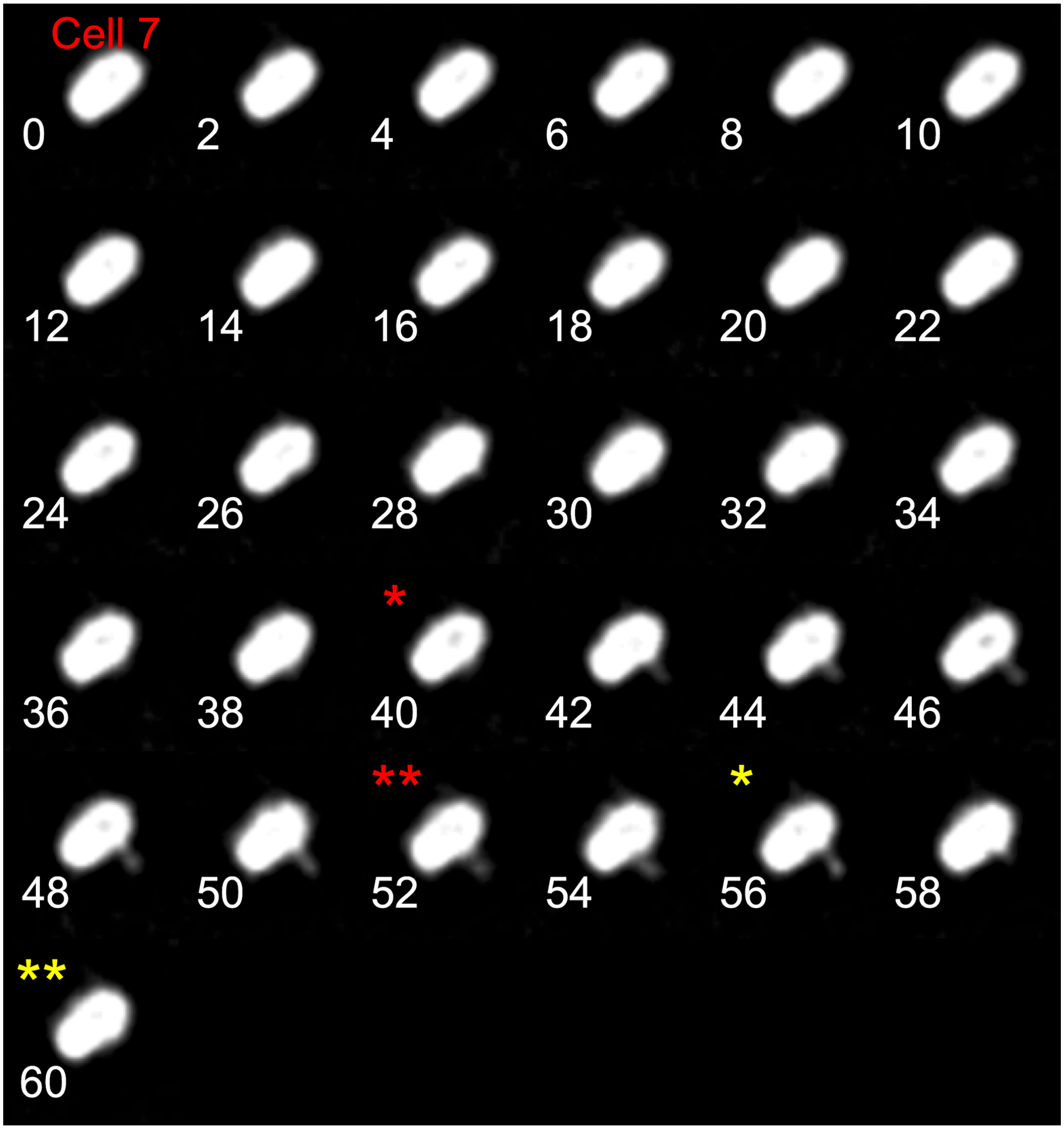


**Figure S13A. Montages of time-lapse imaging of the *ComGC*-Cys cell that was used in Fig. 4 C2 (Cell 7).** Cell were labeled in accord to **Fig. S1**. Time-lapse images were acquired for 1-minute at 2-second intervals, with exposures of 150 ms at a light intensity of 5% for the fluorescent filters, and 15 ms and 32% of exposure and light intensity for DIC. Red * indicates extrusion starting time and red ** indicates extrusion end time. Yellow * indicates retraction starting time and ** indicates for retraction end time.

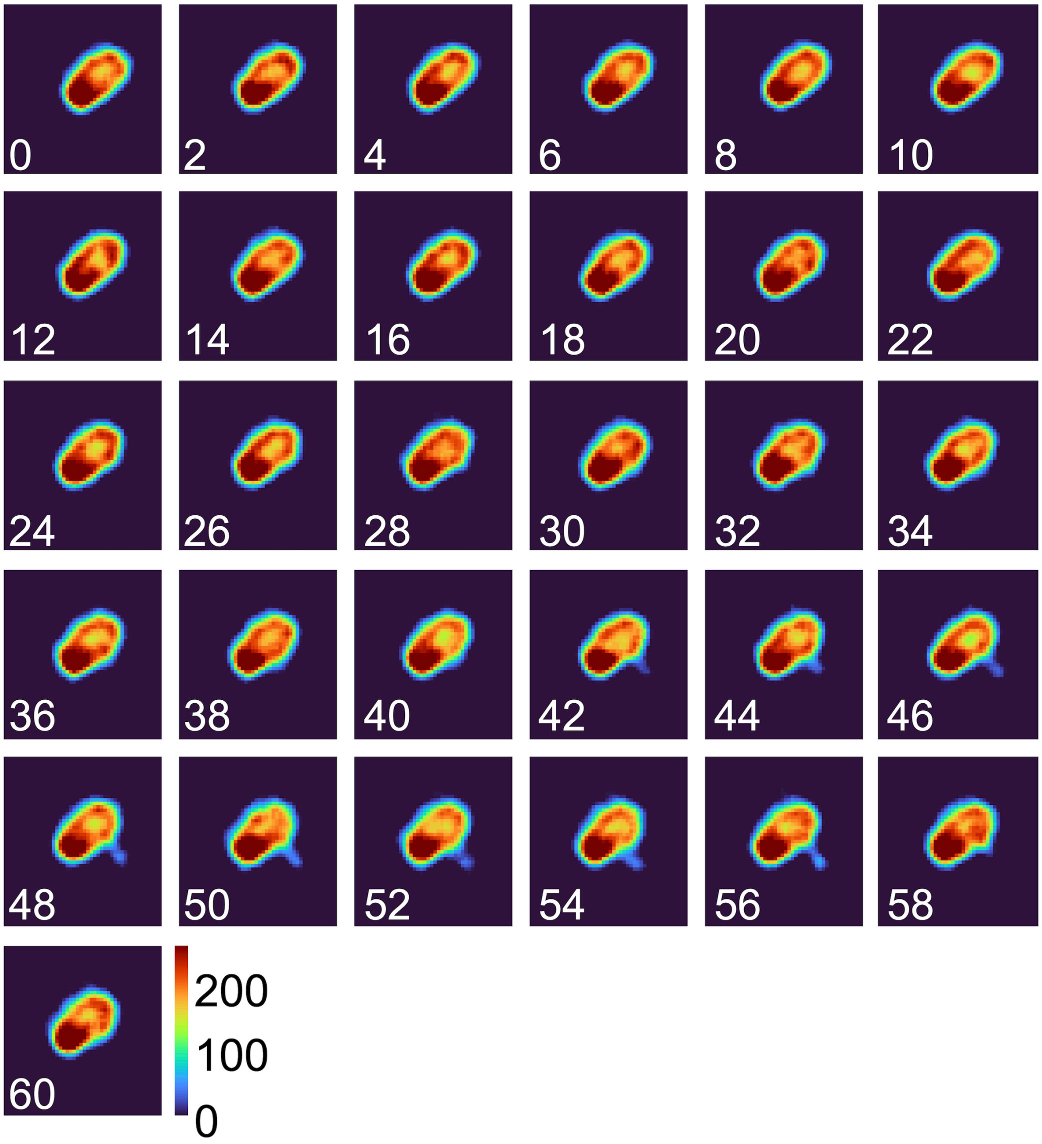


**Figure S13B. Colormap montage of time-lapse images of the *ComGC*-Cys cell that was used in Fig. 4 C2 (Cell 7).** Colormap maps were constructed in MATLAB using 16-bit deconvolved images (scale 0-255).

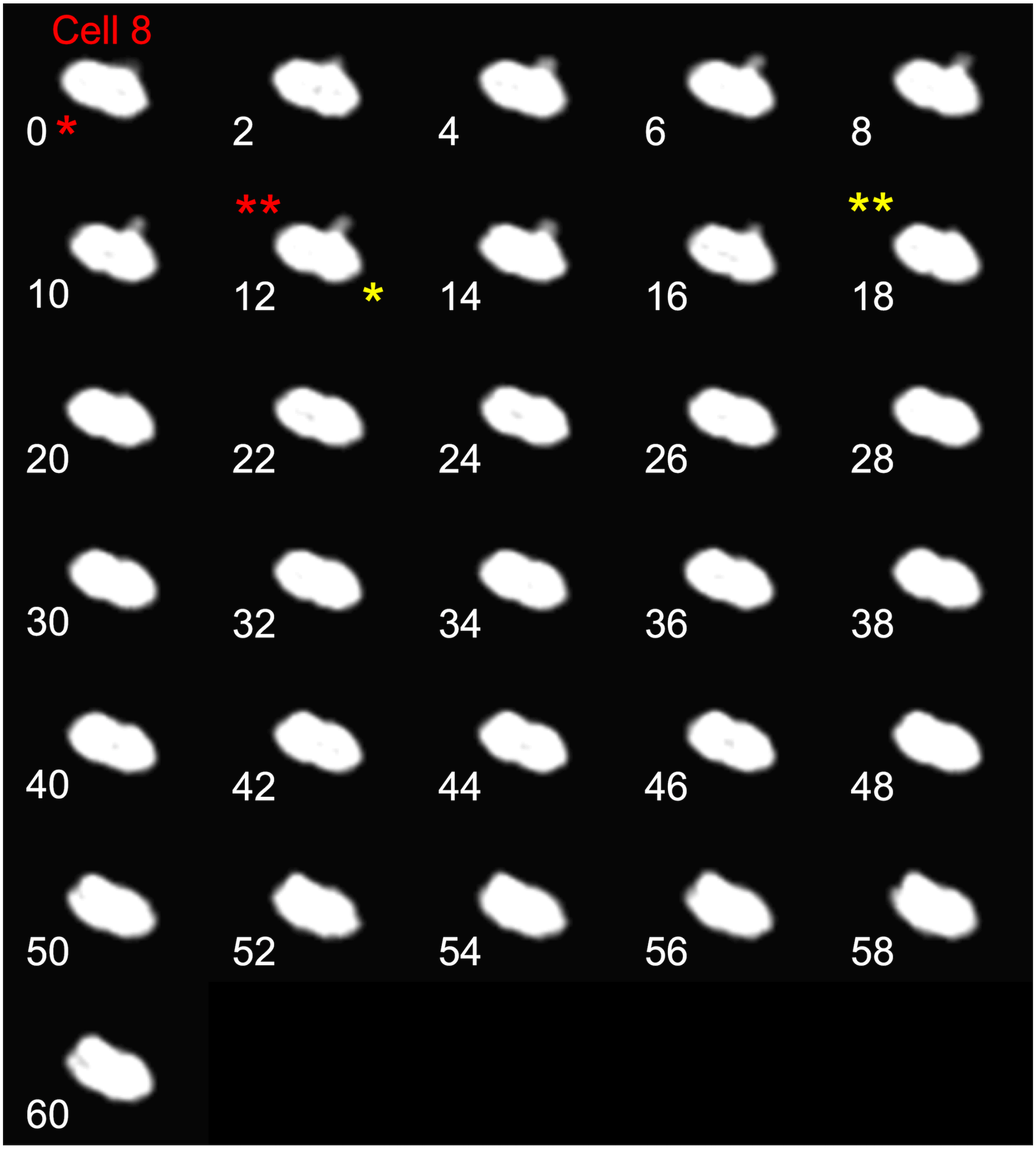


**Figure S14A. Montages of time-lapse imaging of the *ComGC*-Cys cell that was used in Fig. 4 C2 (Cell 8).** Cell were labeled in accord to **Fig. S1**. Time-lapse images were acquired for 1-minute at 2-second intervals, with exposures of 150 ms at a light intensity of 5% for the fluorescent filters, and 15 ms and 32% of exposure and light intensity for DIC. Red * indicates extrusion starting time and red ** indicates extrusion end time. Yellow * indicates retraction starting time and ** indicates for retraction end time.

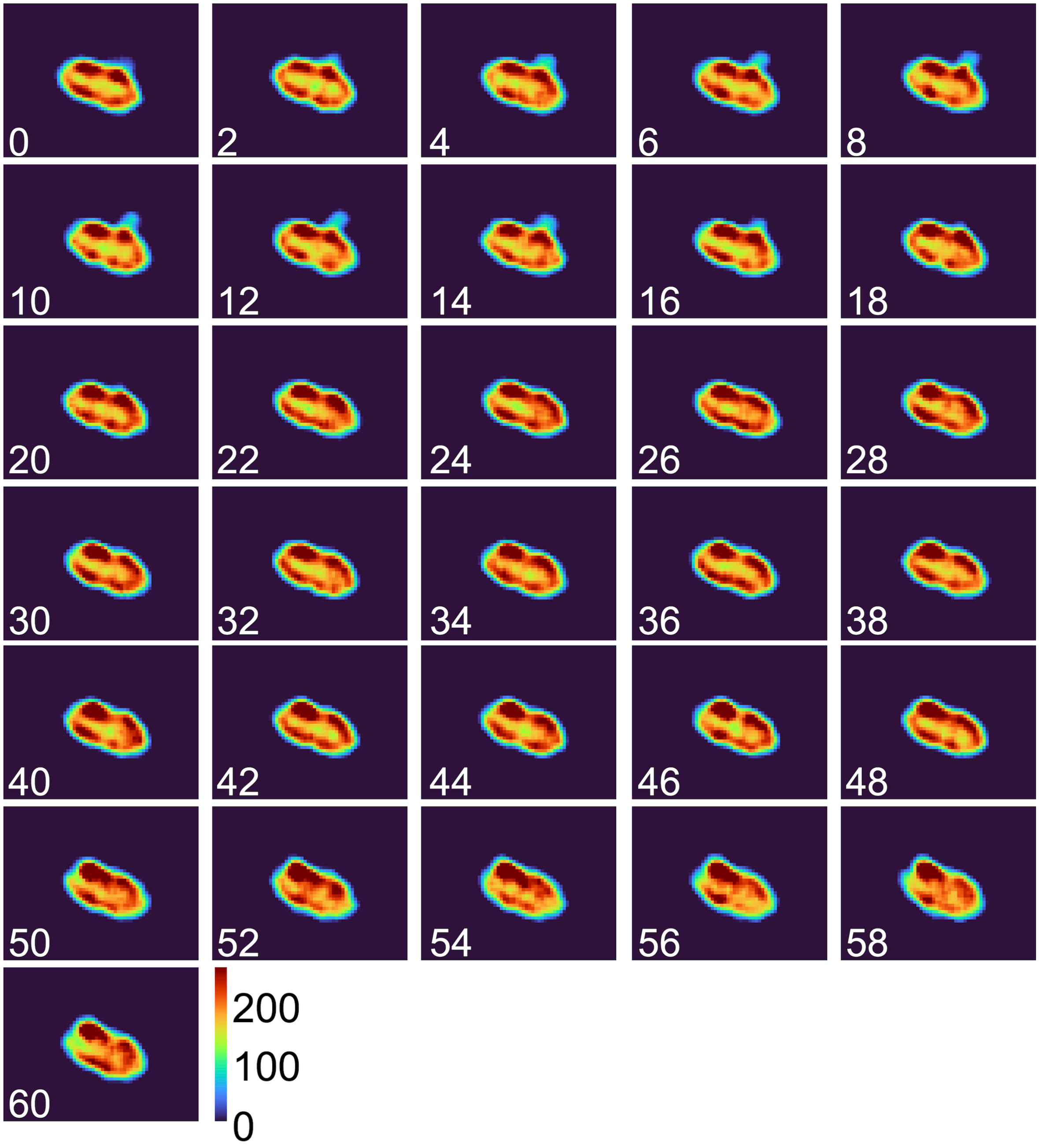


**Figure S14B. Colormap montage of time-lapse images of the *ComGC*-Cys cell that was used in Fig. 4 C2 (Cell 8).** Colormap maps were constructed in MATLAB using 16-bit deconvolved images (scale 0-255).

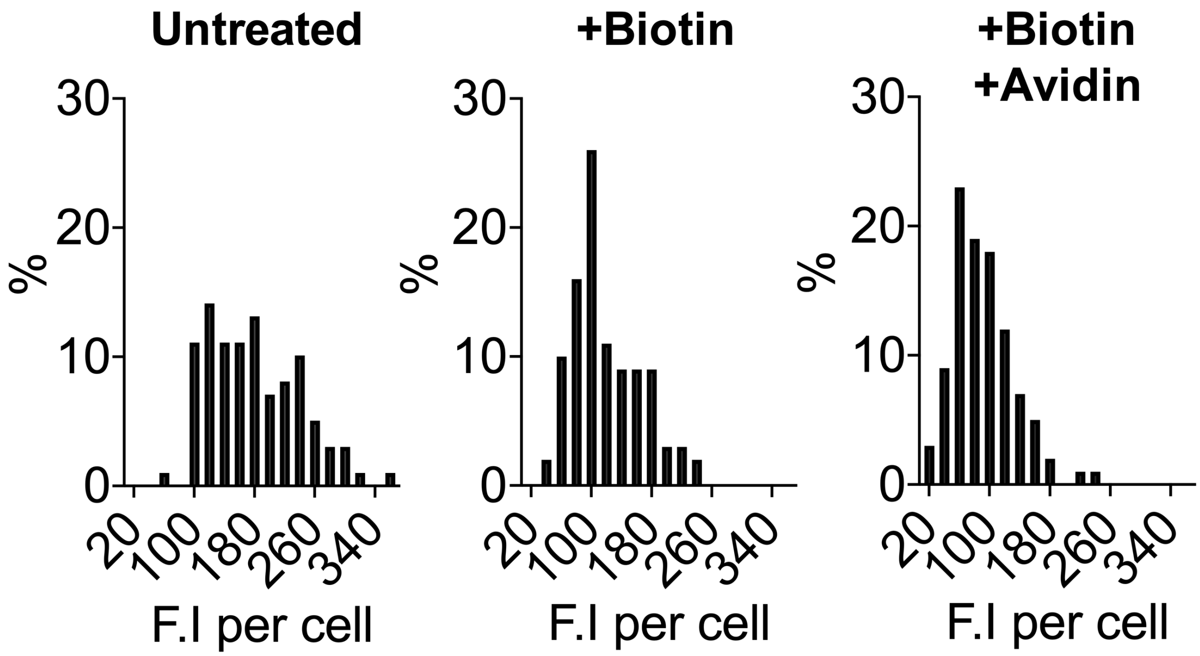


**Figure S15. Fluorescence intensity distributions (N=100 for each condition) of *ComGC*-Cys cells labeled with AF488-mal, biotin-mal plus AF488-mal, with and without Neutravidin.** Cell body was imaged using DIC (32% light intensity, 15 ms exposure) and fluorescent cells and pilus were imaged using a FITC filter (10% light intensity and 200 ms exposure).

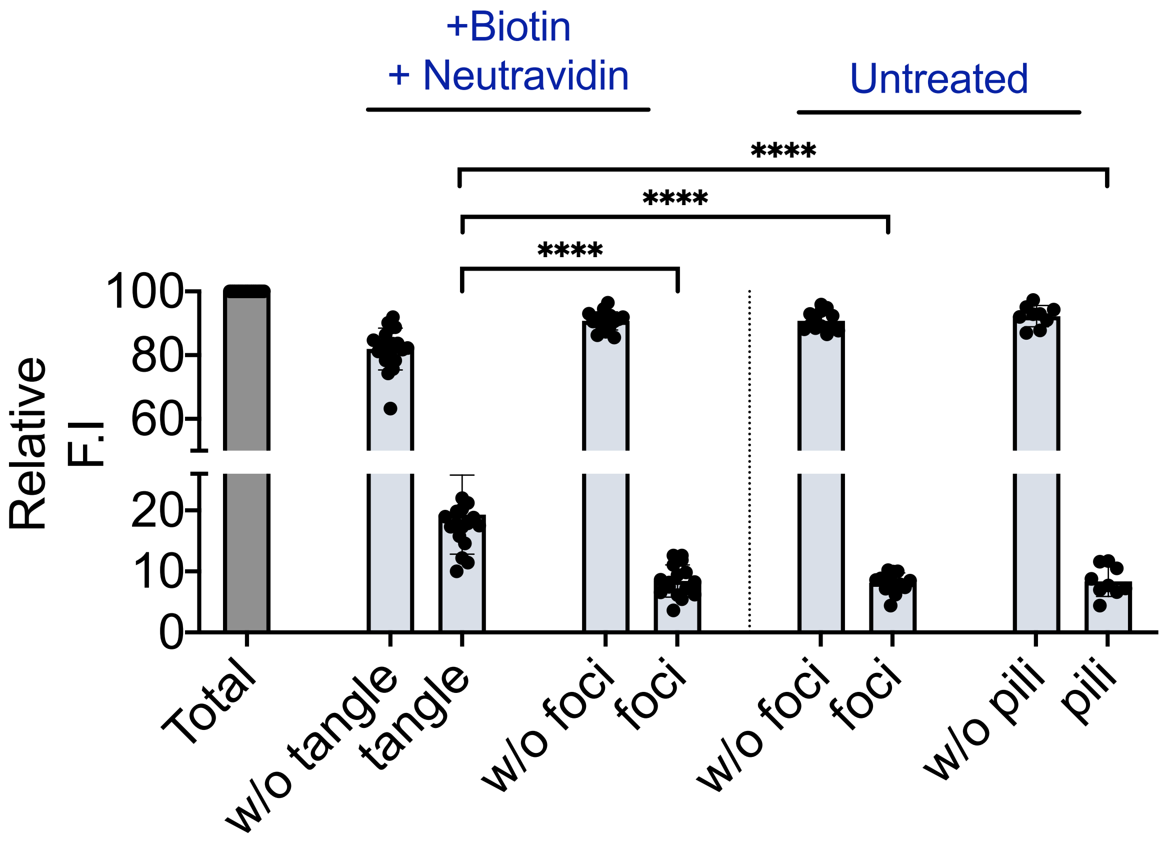


**Figure S16. Comparison of fluorescent intensity of total cell body vs. structures found in *ComGC*-Cys cells treated with only AF488-mal (untreated) or with 5:1 biotin:AF488-mal and neutravidin (+biotin +neutravidin).** Cells with the indicated structure were analyzed with and without the structure, then the signal of the structure was analyzed separately. The integrated fluorescent intensity of cell body including the structure was normalized to 100% (total).

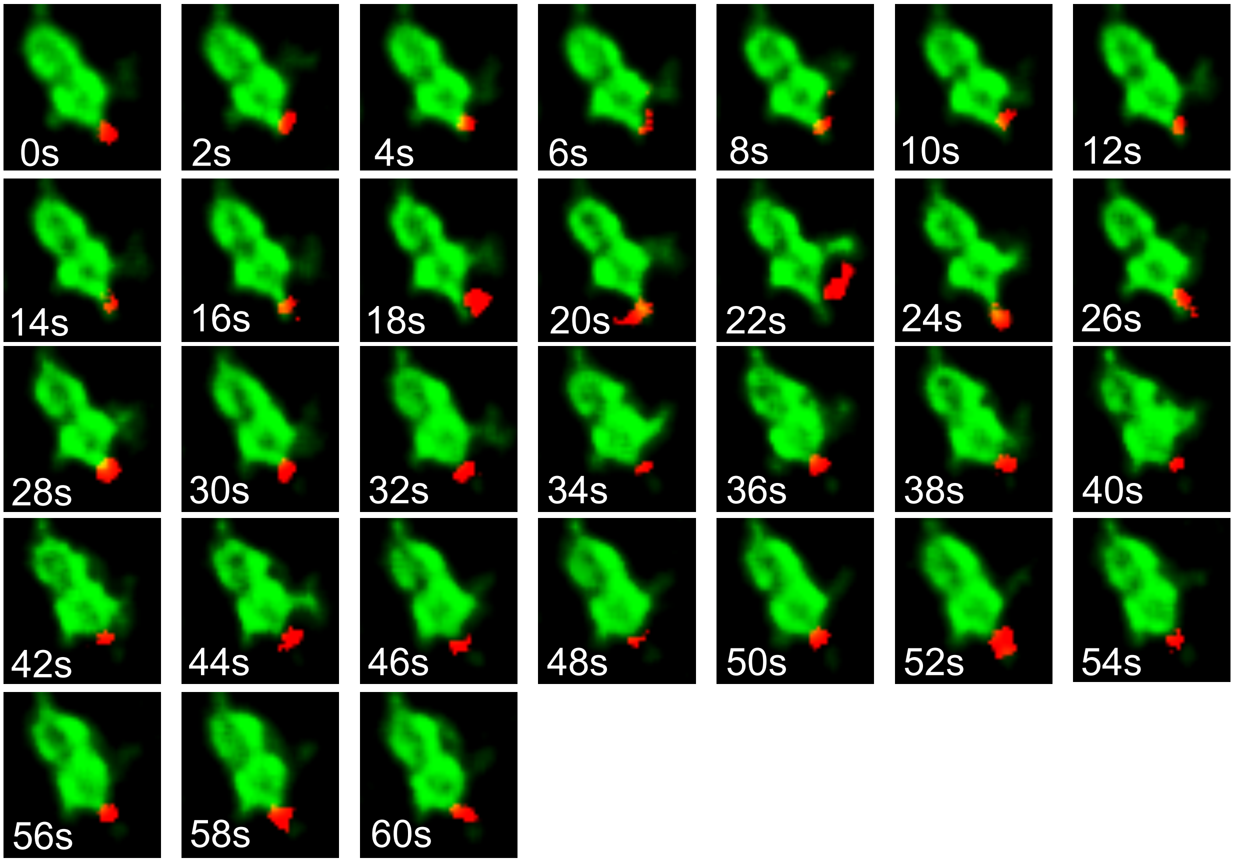


**Figure S17. Full montage of timelapse imaging of interaction of competent *ComGC*-Cys cells (green) with Cy5-lambda DNA (red) shown in Fig. 6.** AF488-labeled cells were mixed 3:1 with 8 μg/mL Cy5-labeled Lambda DNA before applying to a 1% agarose pad for imaging. Cell body and pilus were imaged with DIC phase and FITC filter while Cy5 filter was used for imaging lambda DNA.

**Supplementary video S1:** *ComGC*-Cys cells with AF488-mal labeled mobile pilus

**Supplementary video S2:** *ComGC*-Cys cells with AF488-mal labeled mobile pilus

**Supplementary video S3:** *ComGC*-Cys cells with AF488-mal labeled mobile pilus

**Supplementary video S4:** *ComGC*-Cys cells with AF488-mal labeled mobile pilus

**Supplementary video S5:** Static extrusion of *ComGC*-Cys cells with AF488-mal labeled mobile pilus

**Supplementary video S6:** Static extrusion of *ComGC*-Cys cells with AF488-mal labeled mobile pilus

**Supplementary video S7:** Static extrusion of *ComGC*-Cys cells with AF488-mal labeled mobile pilus

**Supplementary video S8:** S66C *ComGC*-Cys cell extending and retracting labeled pilus (Figure 4 C2, Cell 1)

**Supplementary video S9:** S66C *ComGC*-Cys cell extending and retracting labeled pilus (Figure 4 C2, Cell 2)

**Supplementary video S10:** S66C *ComGC*-Cys cell extending and retracting labeled pilus (Figure 4 C2, Cell 3)

**Supplementary video S11:** S66C *ComGC*-Cys cell extending and retracting labeled pilus (Figure 4 C2, Cell 4)

**Supplementary video S12:** S66C *ComGC*-Cys cell extending and retracting labeled pilus (Figure 4 C2, Cell 5)

**Supplementary video S13:** S66C *ComGC*-Cys cell extending and retracting labeled pilus (Figure 4 C2, Cell 6)

**Supplementary video S14:** S66C *ComGC*-Cys cell extending and retracting labeled pilus (Figure 4 C2, Cell 7)

**Supplementary video S15:** S66C *ComGC*-Cys cell extending and retracting labeled pilus (Figure 4 C2, Cell 8)

**Supplementary video S16:** S66C *ComGC*-Cys cell interaction with Cy5-labeled Lambda DNA.

**Supplementary video S17:** S66C *ComGC*-Cys cell interaction with Cy5-labeled Lambda DNA.

**Supplementary video S18**: S66C *ComGC*-Cys cells and Cy5-labeled Lambda DNA imaging on 1% CDM agarose pads.

**Supplementary video S19**: S66C *ComGC*-Cys cells and Cy5-labeled Lambda DNA imaging on 1% CDM agarose pads.
